# Sex-dependent effects of chronic intermittent voluntary alcohol consumption on attentional, not motivational, measures during probabilistic learning and reversal

**DOI:** 10.1101/2020.05.01.071092

**Authors:** Claudia G. Aguirre, Alexandra Stolyarova, Kanak Das, Sriya Kolli, Vincent Marty, Lara Ray, Igor Spigelman, Alicia Izquierdo

**Author notes:** **Corresponding author:** (AI), or, (CGA).

## Abstract

**Background:** Forced alcohol (ethanol, EtOH) exposure has been shown to cause significant impairments on reversal learning, a widely-used assay of cognitive flexibility, specifically on fully-predictive, deterministic versions of this task. However, previous studies have not adequately considered voluntary EtOH consumption and sex effects on probabilistic reversal learning. The present study aimed to fill this gap in the literature.

**Methods:** Male and female Long-Evans rats underwent either 10 weeks of voluntary intermittent 20% EtOH access or water only (H2O) access. Rats were then pretrained to initiate trials and learn stimulus-reward associations via touchscreen response, and subsequently required to select between two visual stimuli, rewarded with probability 0.70 or 0.30. In the final phase, reinforcement contingencies were reversed.

**Results:** We found significant sex differences on several EtOH-drinking variables, with females reaching a higher maximum EtOH consumption, exhibiting more high-drinking days, and escalating their EtOH at a quicker rate compared to males. During early abstinence, EtOH drinkers made more initiation omissions and were slower to initiate trials than H2O drinking controls, particularly during pretraining. EtOH drinking females were most affected. A similar pattern in trial initiations was also observed in discrimination, but not in reversal learning. EtOH drinking rats were unaffected in their reward collection and stimulus response times, indicating intact motivation and motor responding. Although there were sex differences in discrimination and reversal phases, performance improved over time. We also observed sex-independent drinking group differences in win-stay and lose-shift strategies specific to the reversal phase.

**Conclusions:** Females exhibit increased vulnerability to EtOH effects in early learning: there were sex-dependent EtOH effects on attentional measures during pretraining and discrimination phases. We also found sex-independent EtOH effects on exploration strategies during reversal. Future studies should aim to uncover the neural mechanisms for changes in attention and exploration in both acute and prolonged EtOH withdrawal.

## Introduction

Individuals with Alcohol Use Disorder (AUD) show cognitive impairments, particularly in the domain of cognitive flexibility, broadly defined as the ability to adjust one’s behavior in response to changes in the environment (1). Preclinical models that mimic alcohol (ethanol, EtOH) consumption in humans can elucidate underlying mechanisms related to such cognitive impairments in individuals with AUD (2).

Several groups have tested the effect of forced EtOH exposure on reversal learning, a robust measure of cognitive flexibility commonly used in experimental animals (3) that requires the remapping of reward contingencies. In deterministic (fully-predictive) reversal learning paradigms more frequently employed in behavioral pharmacology experiments, subjects first learn to discriminate and choose between two stimuli, one of which is rewarded and the other which is not. After successful discrimination, the associated outcomes of the two stimuli are reversed, forcing the subject to remap the associations (3–6). In probabilistic reversal learning (PRL) paradigms, each stimulus is associated with a probability of reward (e.g. .80/.20, .70/.30), with one stimulus associated with a higher probability of reward (the “better” option), and another with a lower probability of reward (the “worse” option) (3,7,8).

Voluntary alcohol consumption models (e.g. 2-Bottle Choice, EtOH gelatin) have been used to assess effects on deterministic reversal learning (6,9,10), resulting in no significant treatment group differences. However, it is worth noting that these studies only examined overall performance (i.e. trials to reach criterion, number of correct choices and errors) and did not report whether there were group differences on latency measures to initiate and commit trials and collect reward, or the types of strategies employed on a more fine-grained, trial-by-trial basis. Latency measures, for example, could be used to dissociate processing speed and decision-making from motivational effects. Latency to collect reward along with number of initiated trials are commonly used as a measure of motivation, whereas response latency is often used as a measure of processing or decision-making speed in the Five-Choice Serial Reaction Time Task (5CSRTT) (11–16). Measures of attention vary, but can include number of omissions, as well as percentage of correct responses (i.e. accuracy) (11,13,14,17–19). Although omissions are thought to be indicative of inattentiveness, one must consider this measure in conjunction with response and reward latencies to rule out processing speed and motivational deficits, respectively, as potential confounds (19,20).

We also studied Win-Stay/Lose-Shift (WSLS) strategies commonly analyzed in PRL paradigms. WSLS strategies reflect an animal’s tendency to select the same stimulus after being rewarded (i.e. Win-Stay) or switch to select a different stimulus after not being rewarded (i.e. Lose-shift) (21). Conversely, animals may use less advantageous strategies, such as selecting a different stimulus after being rewarded (i.e. Win-Shift), or choosing the same stimulus after not being rewarded (i.e. Lose-Stay). We probed these in the present study. Also noteworthy, most studies have exclusively used male animals, limiting the generalizability of the results given recent findings showing sex differences in consumption patterns, with female rodents showing higher EtOH intake levels and preference for EtOH (22–26), even exhibiting less aversion to EtOH compared to males (27–30).

The present study sought to address these gaps in the literature by probing the effects of a chronic intermittent voluntary alcohol consumption model on PRL in male and female rats. Rats were administered a 2-bottle choice procedure, during which they were given access to either both 20% EtOH and H2O, or H2O only, 3 days per week for a total of 10 weeks. Five days after their last day of EtOH access they underwent pretraining and advanced to PRL after meeting several training criteria. Given recent findings on sex differences in EtOH consumption, we hypothesized that females would be more EtOH-preferring and reach higher EtOH consumption levels than males. Here, we corroborate previous findings of enhanced EtOH consumption and escalation in female rats compared to male rats. Surprisingly, we found sex-dependent treatment group differences in trial initiations, with the most robust effects in pretraining that carried through early discrimination. These results were in contrast to intact committed trials (i.e., stimulus response) and reward collection times throughout learning. We found changes in WSLS strategy specific to the reversal phase, with EtOH drinkers displaying more exploration (i.e. shift) strategies than H2O drinkers. These effects, were not sex-dependent. Taken together, the present results suggest an enduring effect on attention and exploration-based strategies, but not motivational measures, in stimulus-reward association learning in prolonged abstinence from EtOH.

## Methods

A timeline of all procedures is shown in **Figure 1A**. Subjects were adult male (n=16) and female (n=16) Long-Evans rats (Charles River Laboratories). All animals were between PND 60-70 upon start of EtOH or H2O-only consumption and between PND 130-140 at the start of behavioral testing. All rats underwent a 3 day acclimation period, during which they were pair-housed and given food and water *ad libitum,* and remained in cages with no investigator interference. Following the 3-day acclimation period, animals were handled for 10 min per animal for 5 consecutive days. During the handling period, the animals were given unlimited food and water access and were tail marked. After the handling period, animals were singly-housed under standard housing conditions (room temperature 22–24° C) with a standard 12 h light/dark cycle (lights on at 6am). This study was conducted in strict accordance with the recommendations in the Guide for the Care and Use of Laboratory Animals of the National Institutes of Health. The protocol was approved by the Chancellor’s Animal Research Committee at the University of California, Los Angeles.

**Fig 1.**
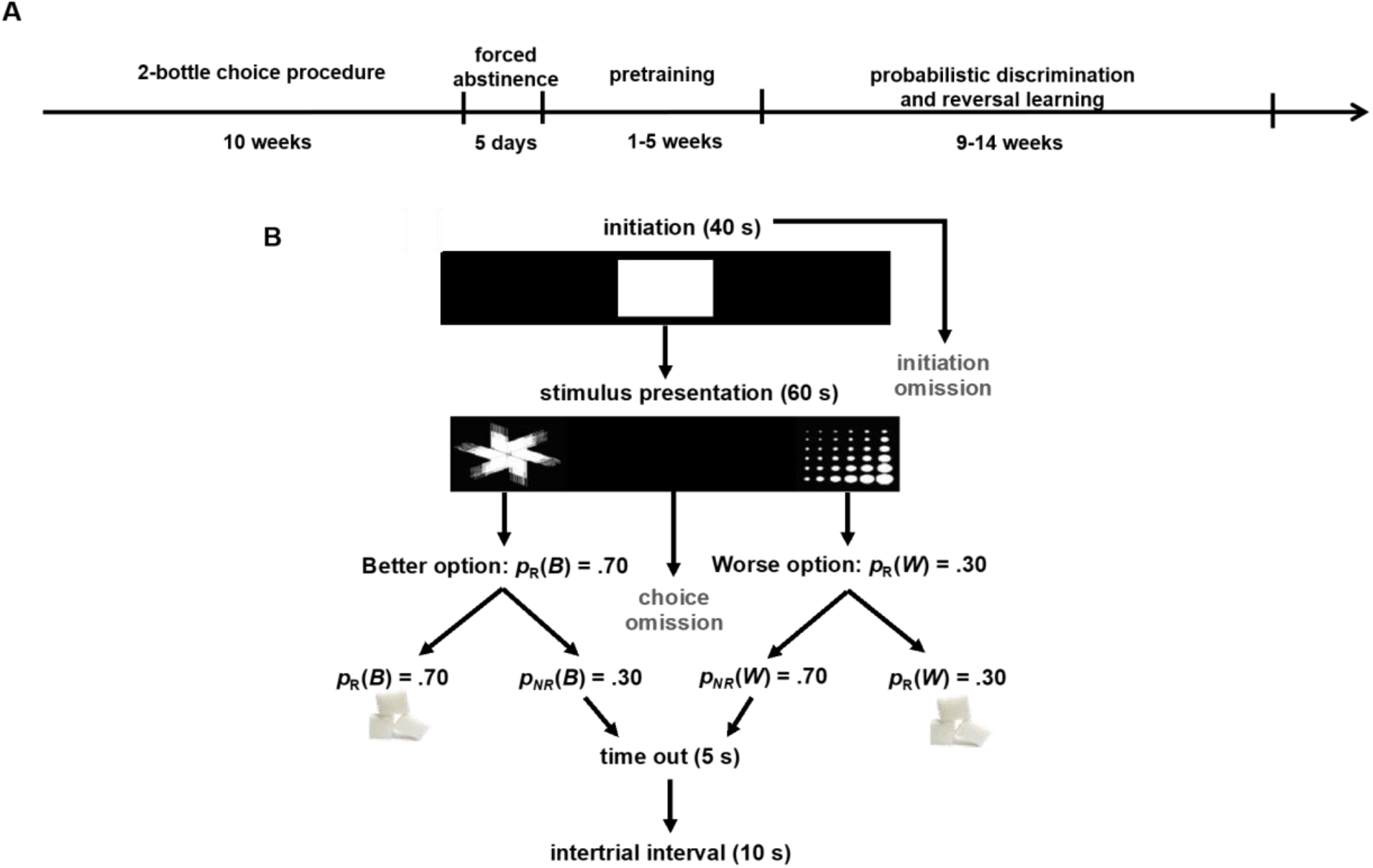
Experimental timeline and probabilistic discrimination and reversal learning paradigm. **(A)** Sequence of events during the voluntary EtOH drinking regimen (i.e. 2-bottle choice procedure), forced abstinence, and behavioral testing, depicted from left to right. **(B)** Task structure of probabilistic reversal learning (PRL), in which the animal initiated a trial by nosepoking the white graphic stimulus in the center screen (displayed for 40 s), and chose between two visual stimuli pseudorandomly presented on either the left and right side of the screen (displayed for 60 s), assigned as the Better or Worse options, rewarded with a sucrose pellet with probability p_R_(B)=0.70 and p_R_(W)=0.30, respectively. If a trial was not rewarded [p_NR_(B) or p_NR_(W)], a 5 s time-out would commence. If a stimulus was not chosen, it was considered an omission and a 10 s ITI would commence. If a trial was rewarded, a 10 s ITI would follow reward collection.

### Rodent voluntary alcohol regimen: 2-bottle choice procedure

Rat home cages were modified to allow for the placement of 2 bottles. Rats (n=16; n=8 male, n=8 female) were given access to both water and 20% alcohol simultaneously, with placement of bottles counterbalanced, for a 24-hour period 3 days per week, and only water on the remaining days. Alcohol access was terminated at 10 weeks (29 days of access) after animals’ EtOH consumption stopped escalating. An age-matched control cohort of water (H2O)-only drinking animals (n=16; n=8 male, n=8 female) was placed in modified home cages allowing for the placement of 2 bottles with water-only also for a total of 10 weeks. Weight of bottles was measured before and after alcohol and/or water-only access to measure daily consumption amounts with a control cage placed on the same rack to account for leakage.

### Behavioral task: rodent probabilistic reversal learning (PRL) task

Immediately following the termination of the consumption period, all animals were placed on food restriction to 14 grams/day (females) or 18 grams/day (males) of chow for 5 days prior to behavioral testing. Animals were weighed every other day and monitored closely to not fall below 85% of their maximum, free-feeding weight. Behavioral testing was conducted in operant conditioning chambers outfitted with an LCD touchscreen opposing the sugar pellet dispenser (31,32) during the animals’ inactive phase. All chamber equipment was controlled by customized ABET II TOUCH software. Following 5 days of forced abstinence (n=16; EtOH group) or a rest period (n=16; H2O group), animals began pretraining.

The pretraining protocol, adapted from established procedures described in Stolyarova et al. (2017) (31), consisted of a series of phases: Habituation, Initiation Touch to Center Training (ITCT), Immediate Reward Training (IMT), designed to train rats to nosepoke, initiate a trial, and select a stimulus to obtain reward. During habituation, rats were required to eat five pellets out of the pellet dispenser inside the chambers within 15 min before exposure to any stimuli on the touchscreen. ITCT began with the display of white graphic stimuli on the black background of the touchscreen. During this stage, a trial could be terminated for one of two reasons: if a rat touched the displayed image (and received reward), or if the image display time (40 s) ended, after which the stimulus disappeared, a black background was displayed, and a 10 s inter-trial interval (ITI) ensued. If the rat did not touch within 40 s this was scored as an *initiation omission.* IMT began in the same way as ITCT, but the disappearance of the white graphic stimulus was now paired with the onset of a target image immediately to the left or right of the stimulus (i.e. forced-choice) that the rat was required to nosepoke to obtain reward. During this stage, a trial could be terminated for one of three reasons. First, if a rat touched the center display (i.e. white graphic stimulus) and touched the image displayed on either side, after which there was a dispensation of one sucrose pellet and illumination of the tray-light. Second, if the rat failed to touch the center white graphic stimulus before the display time ended (40 s), the stimulus disappeared, a black background was displayed, and a 10 s ITI ensued, scored as a *initiation omission.* Third, if the image display time (60 s) ended, after which the stimulus disappeared, a black background was displayed, and a 10 s ITI ensued, scored as a *choice omission*. Rats could also fail to respond to the center stimulus within 40 s during this phase (i.e. initiation omission, as in the previous phase). For habituation pretraining, the criterion for advancement was collection of all 5 sucrose pellets. For ITCT, the criterion to the next stage was set to 60 rewards consumed in 45 min. The criterion for IMT was set to 60 rewards consumed in 45 min across two consecutive days.

After completion of all pretraining schedules, rats were advanced to the discrimination phase of the PRL task, in which they would initiate a trial by touching the white graphic stimulus in the center screen (displayed for 40 s), and choose between two visual stimuli presented on the left and right side of the screen (displayed for 60 s) counterbalanced between trials, assigned as the Better or Worse options, rewarded with a sucrose pellet, with probability pR(B)=0.70 and p_R_(W)=0.30, respectively. Assignment of the stimulus to better or worse reinforcement was counterbalanced across conditions. If a trial was not initiated within 40 s, it was scored as an initiation omission. If a stimulus was not chosen, it was scored as a choice omission, and a 10 s ITI ensued. If a trial was not rewarded, a 5 s time-out would follow, subsequently followed by a 10 s ITI. Finally, if a trial was rewarded, a 10 s ITI would follow after the reward was collected (**Fig 1B**). The criterion was set to 60 or more rewards consumed and selection of the better option in 80% of the trials or higher during a 60 min session across two consecutive days. After reaching criterion for the discrimination phase, the rats advanced to the reversal phase beginning on the next session. During the reversal phase, rats were required to remap stimulus-reward contingencies. The criterion for the reversal phase was the same as the discrimination phase.

### Statistical analyses

To test the study hypotheses, a series of mixed-effects General Linear Models (GLM) and ANOVA analyses were conducted using MATLAB (MathWorks, Natick, Massachusetts; Version R2018b) (33) and SPSS (IBM SPSS Statistics, Version 25) (34). MATLAB was also used for graphing.

The EtOH consumption data were analyzed with ANOVAs with sex and drinking-group as between-subject factors, and EtOH consumption days (D1-D29) as a within-subject factor, with EtOH consumption (g/kg) as the primary outcome (35–37). Independent samples t-tests were conducted for EtOH-drinking related varibles, such as maximum EtOH consumption, calculated as the highest amount of EtOH consumption reached over the course of the 29 days of alcohol access averaged by sex, and the number of high-drinking days (i.e. 5+ g/kg/24 hrs) (38).

Learning data (sessions to criterion, percent correct), number of rewards collected, omission, and latency data were analyzed with GLM in MATLAB *(fitglme* function; Statistics and Machine Learning Toolbox; MathWorks, Natick, Massachusetts; Version R2017a). Percent correct, number of rewards, and initiation omissions, were analyzed using GLM across repeated days of testing per animal with drinking group (EtOH vs. H2O) and sex (female vs male) as fixed effects and animal as a random effect. In Figures, we show the first 15-20 days of learning to avoid overweighting performance of increasingly fewer animals at the extremes. Omission (sums) and latency data (medians) included one observation per subject and were also analyzed with a GLM with drinking group and sex as fixed effects and animal as a random effect. All post-hoc tests were Bonferroni-corrected to account for the number of comparisons. Statistical significance was noted when p-values were less than 0.05, p-values between 0.05 and 0.06 are reported as marginally significant.

Trial-by-trial analyses were conducted to investigate potential sex and group differences in WSLS strategies used in the PRL task. WSLS are strategies commonly examined in decision-making tasks involving risk and reward that can reveal changes in sensitivity to surprising outcomes and feedback learning. Each trial was classified as a *win* if an animal received a sugar pellet, and as a *lose* trial if no reward was delivered. We classified decisions as Win-Stay when a rat chose the same stimulus on the subsequent trial after a win and as Lose-Shift when the rat switched to the alternative stimulus after a loss. We also studied less advantageous strategies: we classified selecting a different stimulus after being rewarded as Win-Shift, and choosing the same stimulus after not being rewarded as Lose-Stay. For win-stay: sum(win-stay)/sum(win); for lose-shift: sum(lose-shift)/sum(lose); for win-shift: 1-sum(win-stay)/sum(win); and for lose-stay: 1-sum(lose-shift)/sum(lose). We compared the frequency of using advantageous strategies (i.e., win-stay, lose-shift) vs. less advantageous strategies (i.e., win-shift, lose-stay) by generating an adaptive score: (win-stay + lose-shift) - (win-shift + lose-stay). Ultimately, we compared the use of advantageous (i.e., win-stay, lose-shift) with less advantageous (i.e., win-shift, lose-stay) strategies by drinking group and sex.

## Results

### EtOH consumption

Independent samples t-tests showed that females reached a greater maximum level of EtOH consumption [t(14)=3.46, p=0.004] (**Fig 2A**) and exhibited more high-drinking days (i.e. days of EtOH consumption 5+ g/kg/24 hours) [t(14)=3.00, p=0.01] than males (**Fig 2B**). A repeated-measures ANOVA was used to assess the within-subject effect of EtOH drinking days and the between-subject factor of sex on EtOH consumption (g/kg). There was a within-subject effect of day [F(28, 392)=6.68, p<0.0001], suggesting an escalation of EtOH over the course of 29 EtOH drinking days (**Fig 2C**). A marginally-significant sex*day interaction was found, [F(28,392)= 1.51, p=0.05], with female animals escalating drinking more steeply than males over the 29 days. Although not significant, we found a trend for a main effect of sex [F(1, 14)=3.42, p=0.09]. Conversely, we saw a de-escalation of water (H2O) consumption over the 29 days of alcohol access, [F(28,392)=13.594 p<0.0001; **Fig 2D**], but found no sex*day interaction, [F(28,392)= 0.993, p=0.48], yet a trend for an effect of sex [F(1, 14)=3.25, p=0.09]. It is important to note that H2O-drinking animals reached a significantly higher body weight 413.44±20.87 (M±SEM), compared to EtOH-drinking animals 332.69±24.09, [t(30)=2.53, p=0.02], and that body weight was significantly negatively correlated with maximum EtOH consumption level [*r*(16)= −0.67, p=0.005], and number of high-drinking days [*r*(16)= −0.61, p=0.01]. Overall females exhibited heavier EtOH consumption patterns than males.

**Fig 2.**
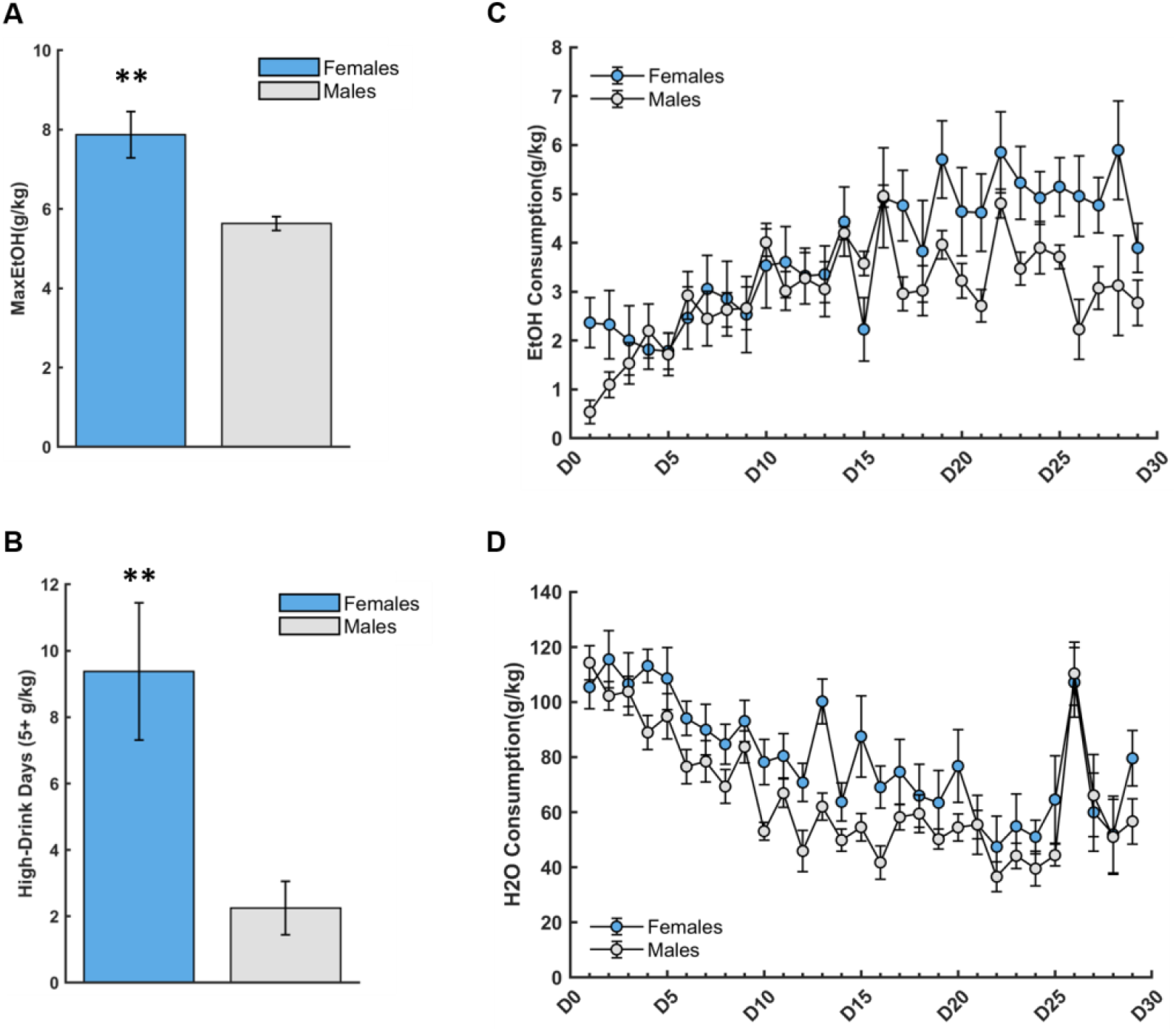
20% Ethanol (EtOH) consumption (g/kg) across drinking days (D1-D29) is more pronounced in females. (**A**) Females reached a greater maximum EtOH level of consumption than males. (**B**) Females exhibited more high-drinking (5-10 g/kg/24 hr) days than males. (**C**) There was a within-subject effect of day, with both males and females escalating their EtOH consumption over time, a marginally significant sex*day interaction, and a trend for a main effect of sex. (**D**) There was a de-escalation of water consumption over the 29 days of alcohol access, but no main effect of sex or sex*day interaction. Bars indicate ± S.E.M. n=8 males, n=8 females, **p<0.01.

### Full Model

Generalized linear models were first run to include all between- (drinking group, sex) and within-subject (learning phase, day) factors, to test for differences among individual factors and (2- and 3-way) interactions on our main behavioral outcomes (i.e. percent correct, denoted as choice of the better option, and rewards collected), omissions (initiation, choice), and latencies (initiation, correct vs. incorrect choice, reward collection), as well as differences in stimulus-based use of advantageous (i.e., win-stay, lose-shift) and less advantageous (i.e., win-lose, lose-stay) strategies.

We found a significant effect of day on choosing the better option (GLM: β_day_ =0.01, p=0.001), indicative of learning over time, as well as a significant group*day interaction (GLM: β_group*day_ =0.02, p=0.01), with the H2O-drinking animals exhibiting higher rates of learning across days. Additionally, there were 3 and 4-way interactions that were significant or marginally-significant, including group*sex*day (GLM: β_group*sex*day_ = −0.03, p=0.01), group*phase*day (GLM: β_group*phase*day_ = −0.02, p=0.05), and group*sex*phase*day (GLM: β_group*sex*phase*day_ =0.02, p=0.03). All animals also increased their number of rewards collected (GLM: β_day_ =1.76, p<.0001), with notable group (GLM: β_group_ =19.70, p=0.02) and sex differences (GLM: β_sex_ =17.26, p=0.04), such that H2O-drinking animals and males collected a greater number of rewards overall compared to EtOH-drinking animals and females, respectively. Significant sex*day (GLM: β_sex*day_= −1.73, p<.0001), and marginally-significant phase*day (GLM: β_phase*day_ = −0.74, p=0.05) interactions indicated that the rate at which rewards were collected increased more rapidly in females and during the discrimination learning phase. We then tested the effect of all factors on initiation omissions, which, as expected, decreased across testing days (GLM: β_day_ = −0.26, p=0.04), but were greater in number among EtOH-drinking animals (GLM: β_group_ = −10.62, p=0.001), females (GLM: β_sex_ = −6.03, p=0.05), and during the discrimination phase (GLM: β_phase_ = −7.42, p=0.02).

Our analyses on omissions resulted in an increased number of initiation omissions in EtOH-drinking animals (GLM: β_group_ = −304.12, p=0.01), and a significant 3-way group*sex*phase interaction for choice omissions (GLM: β_group*sex*phase_ = −3.81, p=0.02). Conversely, H2O-drinking animals exhibited longer reward collection latencies (GLM: β_group_ =0.17, p=0.02), and incorrect choice latencies (GLM: β_group_ = −0.09, p=0.04). Finally, we assessed if there were any differences in use of advantageous vs. less advantageous learning strategies and found a greater use of the advantageous strategies (i.e. win-stay, lose-shift) compared to less advantageous strategies (i.e. win-shift, lose-stay) during the discrimination learning phase (GLM: β_phase_= −0.13, p=0.004). Akaike Information Criteria (AIC) and Bayesian Information Criteria (BIC) were computed for the full model (AIC_percent-correct_— 6375.40; BIC_percent-correct_= −5446.70), as well as for models excluding the phase factor by separately analyzing discrimination (AIC_percent-correct_= −1501.50; BIC_percent-correct_= −1287.40), and reversal learning phases (AIC_percent-correct_= − 1441.2; BIC_percent correct_= − 1234.30), while still including all the other factors (i.e. group, sex, days). The latter models (for which phases were analyzed separately) yielded lower AIC and BIC values, indicative of a better fit. Thus, subsequent analyses were further subdivided into discrimination vs. reversal phases, and early vs. late discrimination and reversal learning based on prior studies indicating these may be particularly informative to contrast (39–44).

### Pretraining (PT) performance

As pretraining was a unique phase with slightly different dependent measures (i.e. forced choice omissions and latencies), a separate GLM was used to assess the effect of group (EtOH, H2O) and sex (female, male), and group*sex interactions on the number of pretraining sessions required to reach criterion to advance to the discrimination learning phase of the PRL task. A significant effect of group and sex emerged, with the EtOH-experienced animals requiring a greater number of pretraining sessions than the H2O-only group (GLM: β_group_ = −14.00, p=0.04; **Fig 3A**), and females requiring a greater number of pretraining sessions than males (GLM: β_sex_ = −11.38, p=0.001; **Fig 3A**). A group*sex interaction was found (GLM: β_group*sex_ =12.00, p=0.01) with EtOH-drinking females requiring more pretraining sessions than EtOH-drinking males (GLM: β_sex_ = −11.38, p=0.03). Both EtOH-drinking females (GLM: β_group_ = −14.00, p=0.01) and EtOH-drinking males (GLM: β_group_ = −2.00, p=0.03) needed more sessions to reach criterion than their H2O-drinking counterparts (**Fig 3A**). There was an overall average of 10.19±1.47 (M±SEM) sessions to successfully meet the pretraining criterion and advance to discrimination learning.

**Fig 3.**
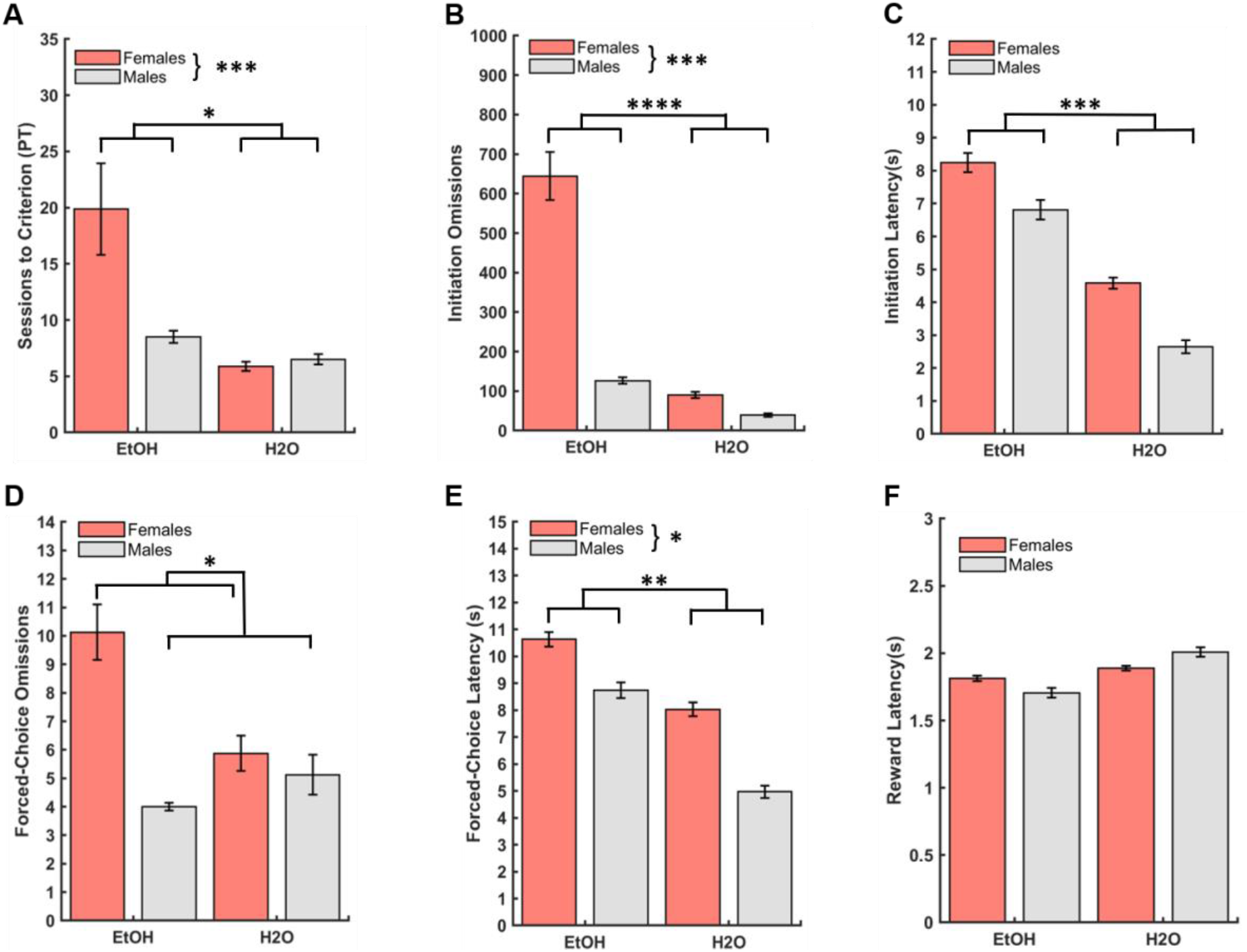
Drinking group, sex differences, and drinking group by sex interactions in operant pretraining. (**A**) The number of pretraining sessions required to reach criterion to advance to the main PRL task was greater for EtOH-drinking (male and female) animals than their H2O-only drinking counterparts, and greater for EtOH-drinking females compared to EtOH-drinking males. Although a significant group*sex interaction was found for this measure, signifcant pairwise comparisons are not depicted for clarity. (**B**) EtOH group and females exhibited more initiation omissions than the H2O group and males, respectively. Although a significant group*sex interaction was also found for this measure, significant pairwise comparisons are not depicted for clarity. (**C**) EtOH group exhibited longer initiation latencies than the H2O group. No sex differences were found for initiation omissions. (**D**) Females displayed more forced-choice omissions. No group differences were found for forced-choice omissions. (**E**) EtOH group and females exhibited longer forced-choice latencies than the H2O group and males, respectively. **(F)** No group or sex differences were found for reward latencies. Latencies represent group medians. Bars indicate ± S. *E. M.* n=16 males, n=16 females, *p ≤0.05, **p ≤0.01, ***p ≤0.001, ****p≤0.0001

We analyzed group and sex differences, as well as the group by sex interaction on initiation omissions, defined as a failure to nosepoke the center white square stimulus within 40s, and initiation latencies, defined as the duration until nosepoke of the center white square stimulus. There was an effect of both group (GLM: β_group_ = −554.12, p=0.0001; **Figs 3B**), and sex (GLM: β_sex_ = −517.87, p=0.0003; **Figs 3B**) on initiation omissions, with EtOH-drinking animals and females displaying more initiation omissions compared to H2O-drinking animals and males, respectively. A significant drinking group*sex interaction (GLM: β_group*sex_ = 467.00, p=0.01; **Fig 3B**), indicated that both EtOH-drinking females (GLM: β_group_ = −554.00 p=0.01) and males (GLM: β_group_ = −87.13, p=0.01) exhibited more initiation omissions than their H2O-drinking counterparts, and EtOH-drinking females exhibited more initiation omissions than EtOH-drinking males (GLM: β_sex_ = −517.88, p=0.01). The EtOH group also displayed longer initiation latencies than the H2O group (GLM: β_group_ = −3.48, p=0.001; **Figs 3C and S3A**), but there was no effect of sex (GLM: β_sex_ = −1.37, p=0.18; **Figs 3C and S3D**), or group*sex interaction (GLM: β_group*sex_ = −0.09, p=0.95; **Fig 3C**).

Next we analyzed differences in forced-choice omissions, defined as failure to nosepoke the stimulus presented on either the left or right side of the touchscreen after initiation of the trial, and forced-choice latencies, defined as duration until nosepoke of the stimulus presented on the left or right side. There was a significant effect of sex (GLM: β_sex_ = −6.13, p=0.03; **Fig 3D**), with females displaying more forced-choice omissions, but no significant group (GLM: β_group_ = −4.25, p=0.13; **Fig 3D**), or group*sex interaction (GLM: β_group*sex_ = 5.38, p=0.17; **Fig 3D**). There was a significant effect of both group (GLM: β_group_ = −2.76, p=0.01; **Fig 3E and S3B**), and sex (GLM: β_sex_ = −2.37, p=0.03; **Fig 3E and S3E**), on forced-choice latency, with the EtOH-experienced animals and females taking longer to select the stimulus, but no significant group*sex interaction (GLM: β_group*sex_ = −0.45, p=0.76). Finally, we analyzed data for differences in reward latency, defined as the duration to collect the sucrose reward from the food magazine, and found no effect of group (GLM: β_group_ = 0.44, p=0.70; **Figs 3F and S3C**), sex (GLM: β_sex_ = −0.01, p=0.92; **Figs 3F and S3F**), or group*sex (GLM: β_group*sex_ = 0.15, p=0.35; **Fig 3F**) interaction.

Collectively, the omission and latency data for initiations and forced-choice trials suggest an attenuating effect of EtOH experience, and specifically in EtOH- experienced females, on quickly responding to stimuli, while the reward collection data point to preserved motor responding and motivation for reward in EtOH-experienced animals.

### Probabilistic Discrimination (D) performance

There were no significant group (GLM: β_group_ = −9.50, p=0.07) or sex differences (GLM: β_sex_ = 3.13, p=0.54) on total number of sessions to reach criterion for the discrimination phase, but a marginally-significant group*sex interaction on this measure was observed (GLM: β_group*sex_ =14.88, p=0.047; **Fig 4A)**. Overall, all animals performed comparably regardless of group or sex, with an overall average of 27.41±2.17 days required to successfully discriminate and advance to the reversal phase.

**Fig 4.**
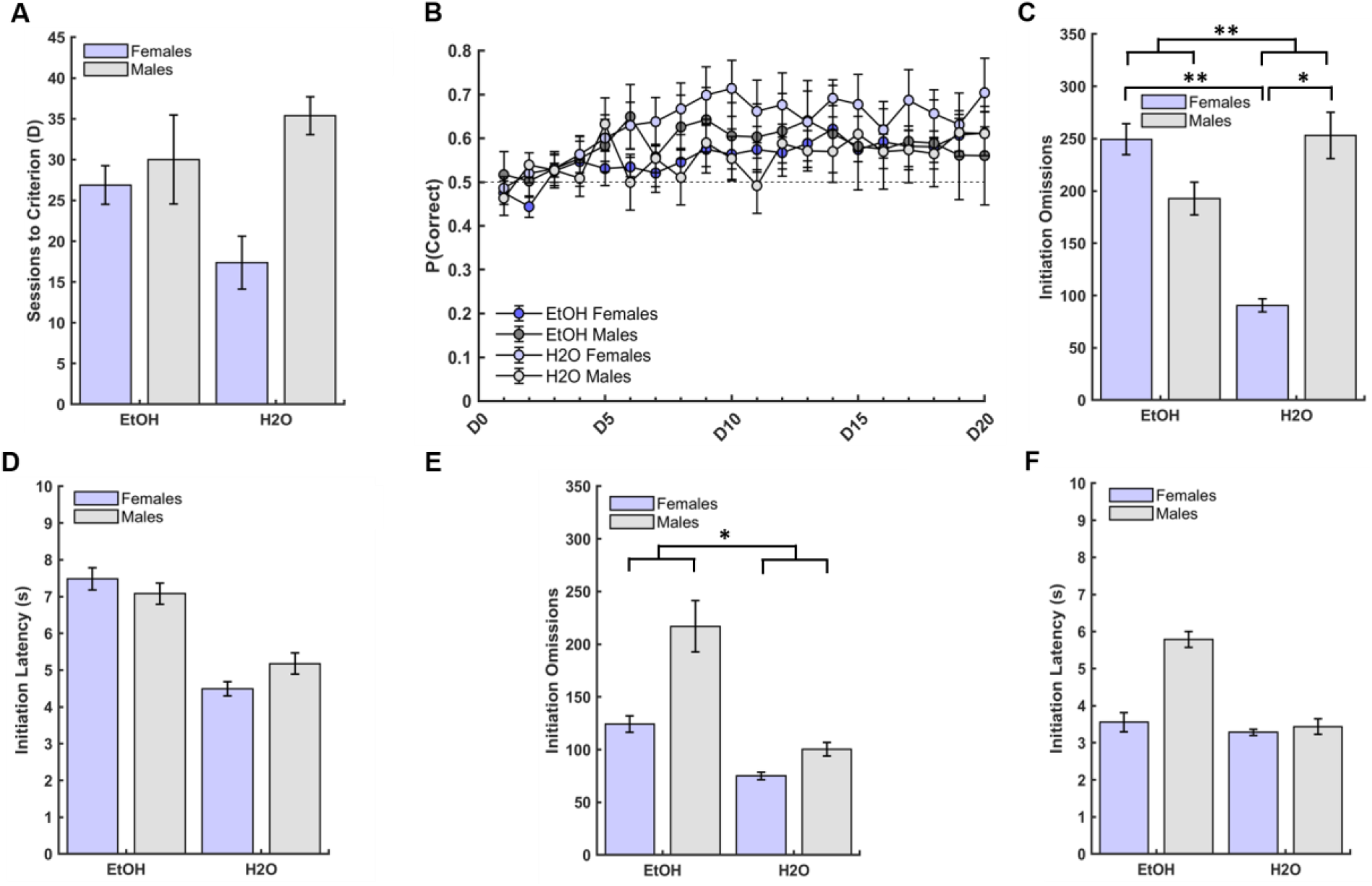
Drinking group differences in initation omissions, but not latencies, during early and late probabilistic discrimination learning performance. (**A**) There were no group or sex differences in the number of sessions to reach criterion for the discrimination learning phase, only a marginally significant group*sex interaction. (**B**) There was no effect of group or sex on probability of choosing the better option, only an effect of day (D1-D20), with the probability of choosing the better option increasing across testing days irrespective of sex or group. (**C**) EtOH group exhibited more initiation omissions compared to the H2O group. EtOH-drinking females exhibited more initiation omissions than H2O-drinking females, and H2O-drinking males exhibited marginally more initiation omissions than H2O-drinking females in early discrimination learning. (**D**) No group or sex differences emerged for initiation latencies during early discrimination learning. (**E**) EtOH-drinking animals (male and female) exhibited marginally more initiation omissions than H2O-drinking animals, but no sex differences emerged for late discrimination learning. (**F**) No group or sex differences were found for initiation latencies during late discrimination learning. Latencies were medians. Bars indicate ± *S. E. M.* n=16 males, n=16 females, *p ≤0.05, **p ≤0.01

A GLM model was used to test the effect of drinking group (EtOH, H2O), sex (female, male), days, and their 2-way and 3-way interactions on percent correct (i.e. choosing the better option), number of rewards (i.e. sucrose pellets), and initiation omissions across 20 testing days of discrimination learning (D1-D20). All animals demonstrated learning by showing an increase in choosing the better option across days, (GLM: β_day_ =0.01, p=0.001; **Figs 4B)**, regardless of group or sex. There was no effect of drinking group (GLM: β_group_ = −0.01, p=0.79), or sex (GLM: β_sex_ =0.07, p=0.17), and no group*sex (GLM: β_group*sex_ = −0.03, p=0.65), or sex*day (GLM: β_sex*day_ =0.01, p=1.00) interaction on percent correct. There was however a significant group*day interaction (GLM: β_group*day_ = 0.02, p=0.02), with the H2O-drinking animals choosing the better option increasingly more across days than the EtOH-drinking animals. There was also a significant group*sex*day interaction (GLM: β_group*sex*day_ = −0.02, p=0.01). Post-hoc comparisons revealed males learned quicker than females (GLM: β_sex_= 0.01, p<0.001), that H2O-drinking females chose the better option increasingly more across days than their EtOH-drinking counterparts (GLM: β_group_ =0. 01 p=0.002); and that the same pattern was not observed for males.

All animals increased the number of rewards collected across days, (GLM: β_day_ =1.83, p=<0.0001; **Fig S1A**). Drinking group (GLM: β_group_=22.59, p=0.01), and sex differences (GLM: β_sex_ =17.60, p=0.04) also emerged, with both H2O-drinking animals and males collecting a greater number of rewards (**Fig S1A**). No significant group*sex (p=0.07), group*day (p=0.78), or group*day*sex (p=0.08) interactions on number of rewards collected were found, only a significant sex*day interaction (GLM: β_sex*day_ = −1.75, p<0.0001), with females displaying a greater increase in rewards collected across days (GLM: β_day_ =1.69, p<0.0001), compared to males (**Fig S1A**).

Conversely, all animals decreased the number of initiation omissions across days, (GLM: β_day_ = −0.29, p=0.02; **Fig S1B**), regardless of group or sex. There were significant group (GLM: β_group_ = −11.17, p=0.001) and sex differences (GLM: β_sex_ = −6.36, p=0.04), with EtOH-drinking animals and females exhibiting more initiation omissions than H2O-drinking and males, respectively (**Fig S1B**). A significant group*sex interaction (GLM: β_group*sex_ =14.81, p=0.01), with EtOH-drinking females displaying more initiation omissions than H2O-drinking females (GLM: β_group_ = −304.13, p=0.02); the same pattern was not observed in males (**Fig S1B**). Similarly, a significant group*sex*day (GLM: β_group*sex*day_ = −0.69, p=0.02) revealed that EtOH-drinking females decreased their number of initiation omissions across discrimination learning at a slower rate than EtOH-drinking males (p=0.03) and H2O-drinking females (p=0.03). There were no differences between male and female H2O drinkers, or between EtOH- vs. H2O-drinking males. Finally, we conducted GLM analyses on total omissions and median latencies in discrimination learning. Our results indicated EtOH-drinking animals exhibited more initiation omissions than H2O-drinking animals (GLM: β_group_ = −304.12, p=0.01), whereas H2O-drinking animals had longer reward collection latencies than EtOH-drinking animals (GLM: β_group_=0.17, p=0.02). All other types of omission and latency analyses yielded non-significant results.

#### First 500 trials

Because interesting latency differences were obtained for the pretraining phase and based on prior studies comparing early vs. late discrimination learning, we conducted further latency analyses for other phases of learning to assess if these trends were maintained. An analysis of initiation latencies and omissions, choice (correct and incorrect) latencies and omissions, and reward latencies was conducted for the first 500 trials of discrimination learning to capture early learning in this phase, with animals averaging 89.19±3.38 committed trials per day. There was a significant group difference on initiation omissions (GLM: β_group_ = −158.75, p=0.002; **Fig 4C**), with EtOH-drinking animals exhibiting more initiation omissions than H2O-drinking animals, but no sex differences (GLM: β_sex_ = −56.50, p=0.36; **Fig 4C**). A significant drinking group*sex interaction on initiation omissions emerged (GLM: β_group*sex_ = 219.00, p=0.02; **Fig 4C**), with EtOH-drinking females exhibiting more initiation omissions than their H2O-drinking female counterparts (p=0.01), and H2O-drinking males exhibiting marginally more initiation omissions than H2O-drinking females (p=0.05), but EtOH-drinking males were no different than H2O- drinking males. There was no effect of drinking group (**Figs 4D and S4A**), sex (**Figs 4D and S4E**), or group*sex interaction (**Fig 4D**) on initiation latencies, choice omissions, correct or incorrect choice latencies, or reward collection latencies (**Figs S4B and S4F**), with the exception of a marginally significant group effect (GLM: β_group_ =0.88, p=0.05), with H2O-drinking animals exhibiting more choice omissions than EtOH-drinking animals. It should be noted that choice omissions represented a small number of occurrences (normally ranging from 0-2) and this effect was driven by a single outlier with 4 choice omissions, which upon removal yielded a nonsignificant group effect (GLM: β_group_ =0.48, p=0.14). In summary, the data for early discrimination learning suggest an enduring effect of EtOH experience on initiating trials, with females most affected.

#### Last 500 trials

An analysis of initiation latencies and omissions, choice (correct and incorrect) latencies and omissions, and reward latencies was also conducted for the last 500 trials of discrimination learning to capture late phase learning, with animals averaging 118.98±3.89 committed trials per day. There was a marginally-significant effect of group (GLM: β_group_ = −49.13, p=0.05; **Fig 4E**), with EtOH-drinking animals exhibiting more initiation omissions than H2O- drinking animals, but no sex differences (GLM: β_sex_ = 92.88, p=0.21; **Fig 4E**) or group*sex interaction (GLM: β_group*sex_ = −67.50, p=0.38; **Fig 4E**). There were no group (GLM: β_group_ = −0.92, p=0.24; **Figs 4F and S4C**), or sex differences (GLM: β_sex_ =1.26, p=0.20; **Figs 4F and S4G**) on initiation latencies, as well as no group*sex interaction (GLM: β_group*sex_ = −0.76, p=0.51; **Fig 4F**). We did, however, find a significant effect of group on incorrect choice latencies (GLM: β_group_ = 0.28, p=0.02), with H2O-drinking animals displaying longer latencies than EtOH-drinking animals, but no significant effect of sex, or group*sex interaction was found for this measure. Similar to what was found during early discrimination, there was a significant group effect on choice omissions in late discrimination (GLM: β_group_ =1.13, p=0.01), with the H2O-drinking animals exhibiting more choice omissions than the EtOH-drinking animals, but no effect of sex, or group*sex interaction was found for this measure. However, this effect was largely driven by the same animal as during early discrimination, which upon removal, yielded non-significant results (GLM: β_group_ =0.43, p=0.06). Finally, our results indicated that H2O-drinking animals displayed longer reward collection latencies than EtOH-drinking animals (GLM: β_group_ =0.23, p=0.01; **Fig S4D**), but no effect of sex **(Fig S4H**) or group*sex interaction emerged. Thus, the pattern of prior EtOH experience rendering animals more likely to fail to initiate trials was also observed through late discrimination learning, but rats did not take longer to initiate trials when they did so, as in pretraining and early discrimination phases. Although we did observe greater choice omissions by the H2O-drinking animals during both early and late discrimination, this was driven largely by one animal.

### PRL reversal (R) performance

There were no significant group differences (GLM: β_group_ = 5.75, p=0.18), sex differences (GLM: β_sex_ = −5.25, p=0.22), or group*sex interaction (GLM: β_group*sex_ = −1.88, p=0.74) on sessions to reach criterion for the reversal learning (**Fig 5A)**. There was an overall average of 25.27±1.62 days to successfully complete the PRL phase.

**Fig 5.**
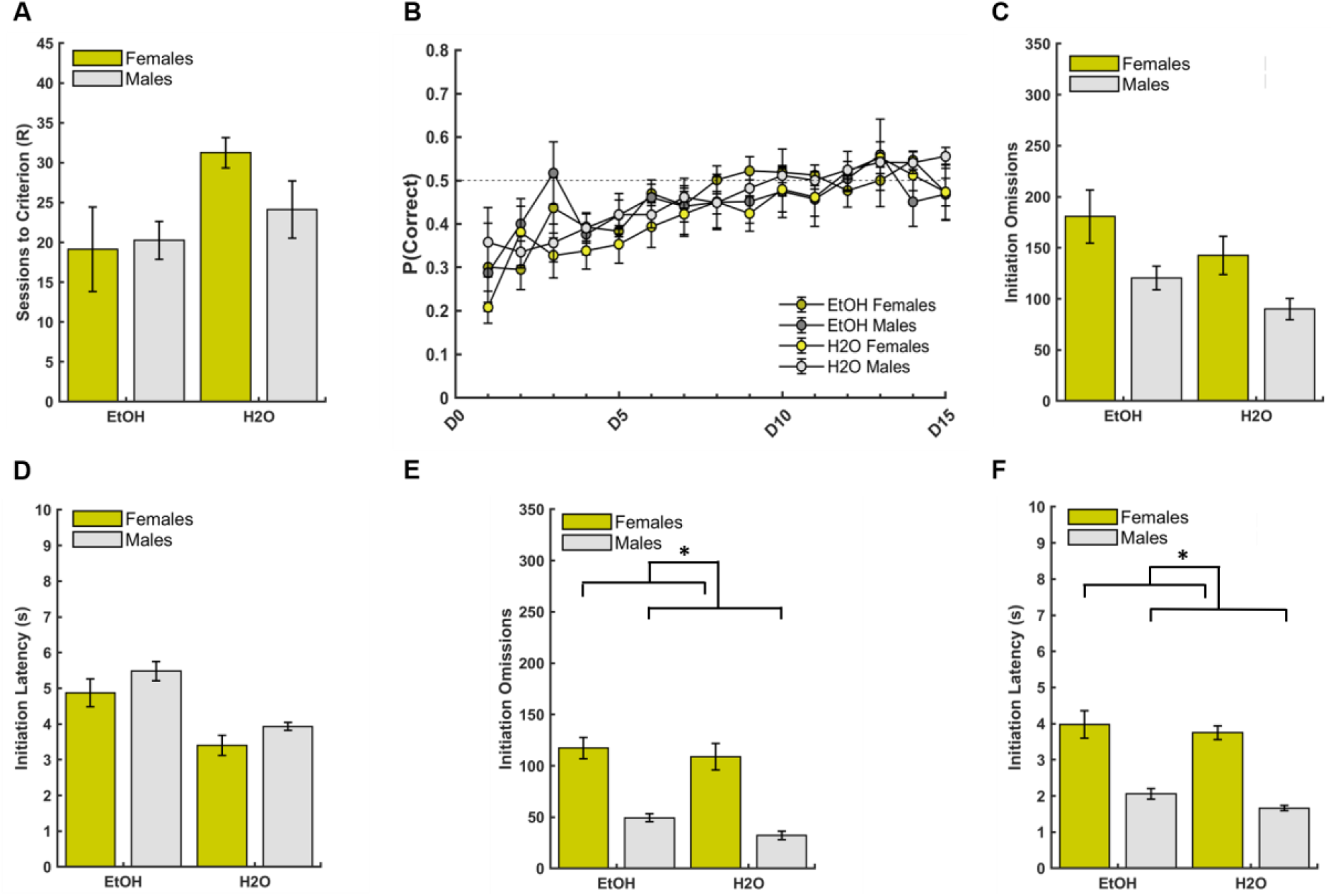
Sex differences, but no drinking group differences, in initiation omissions and latencies during late probabilistic reversal learning performance. (**A**) There were no group or sex differences in the number of sessions to reach criterion for the reversal learning phase. (**B**) There was no effect of group or sex on probability of choosing the better option, only an effect of day (D1-D25), with the probability of choosing the better option increasing across the fifteen testing days regardless of sex or group. (**C**) No group or sex differences were found for initiation omissions during early reversal learning. (**D**) No group or sex differences were found for initiation latencies during early reversal learning. (**E**) Females exhibited more initiation omissions than males during late reversal learning. (**F**) Females exhibited marginally longer initiation latencies than males during late reversal learning. Latencies represent group medians. Bars indicate ± *S. E. M.* n=16 males, n=14 females, *p ≤0.05

A GLM model was used to test the effect of drinking group (EtOH, H2O), sex (female, male), days, and their 2-way and 3-way interactions on probability of choosing the better option, number of rewards, and initiation omissions across 15 days of reversal learning. Two females were excluded because they either failed to meet criterion for discrimination learning, and never advanced to reversal learning. All other animals demonstrated learning by exhibiting an increase in choosing the better options across days, (GLM: β_day_ =0.01, p=0.01; **Fig 5B)**, irrespective of group or sex. There was no effect of group (GLM: β_group_ = −0.07, p=0.10), or sex (GLM: β_sex_ = − 0.04, p=0.51), and no group*sex (GLM: β_group*sex_ = 0.10, p=0.17), sex*day (GLM: β_sex*day_ =0.002, p=0.60), group*day (GLM: β_group*day_ =0.002, p=0.47), or group*sex*day (GLM: β_group*sex*_ day = − 0.004, p=0.46) interactions on the probability of choosing the better option.

All animals increased the number of rewards collected over days, (GLM: β_day_ =0.82, p=0.01; **Fig S1C**). There was no significant effect of group (GLM: β_group_=3.69, p=0.75), or sex (GLM: β_sex_ =2.40, p=0.85), and no significant group*sex (GLM: β_group*sex_ =5.73, p=0.74), group*day (GLM: β_group*day_= −0.02, p=0.95), sex*day (GLM: β_sex*day_=0.66, p=0.08), or group*sex*day (GLM: β_group*sex*day_=0.48, p=0.40) interactions on number of rewards collected. All animals decreased the number of initiation omissions across days, (GLM: β_day_ =0.10, p=0.55; **Fig S1D**), regardless of group or sex. There was was no significant effect of group (GLM: β_group_= −0.44, p=0.91), or sex (GLM: β_sex_ = −1.81, p=0.61), and no significant group*sex (GLM: β_group*sex_ =0.67, p=0.91), group*day (GLM: β_group*day_= −0.06, p=0.78), sex*day (GLM: β_sex*day_=0.09, p=0.70), or group*sex*day (GLM: β_group*sex*day_= −0.51, p=0.09) interactions on initiation omissions. Finally, we conducted GLM analyses on total omissions and median latencies in reversal learning. Similar to the above measure for across- day learning, we found no significant effect of group (GLM: β_group_ =7.88, p=0.97), sex (GLM: β_sex_ = −234.25, p=0.17), or group*sex (GLM: β_group*sex_ = −122.1, p=0.59) interaction for initiation omissions. However, though there was no effect of sex (GLM: β_sex_=0.67, p=0.49), we found that H2O-drinking animals exhibited more choice omissions than EtOH-drinking animals (GLM: β_group_ =2.54, p=0.03), with a significant group*sex interaction revealing H2O-drinking females exhibited more choice omissions than EtOH-drinking females and H2O-drinking males (GLM: β_group*sex_= −3.29, p=0.03). Additionally, there was a marginally-significant effect of group on incorrect choice latencies (GLM: β_group_=0.15, p=0.05), with H2O-drinking animals exhibiting longer incorrect choice latencies than EtOH-drinking animals. Analyses on initiation, incorrect, and reward latencies yielded non-significant results. Unlike for the discrimination phases, upon removal of 2 outliers, the effect of drinking group remained significant, with H2O-drinking animals exhibiting more choice omissions than EtOH-drinking animals (GLM: β_group_=1.59, p=0.03).

#### First 500 trials

An analysis of initiation latencies and omissions, choice (correct and incorrect) latencies and omissions, and reward latencies was conducted for the first 500 trials of reversal learning, to capture early reversal learning. Animals averaged 119.70±8.32 committed trials per day. There were no significant group (GLM: β_group_ = −38.04, p=0.65; **Fig 5C**), or sex differences (GLM: β_sex_ = −60.29, p=0.41; **Fig 5C**) on initiation omissions, and no significant group*sex interaction (GLM: β_group*sex_ = 7.67, p=0.94; **Fig 5C**). Similarly, there was no effect of drinking group (**Figs 5D and S5A**), sex (**Figs 5D and S5E**), or group*sex interaction (**Fig 5D**) on initiation latencies, choice omissions, correct choice latencies, or reward collection latencies (**Figs S5B and S5F**), with the exception of a significant group difference on incorrect choice latencies (GLM: β_group_ =0.32, p=0.004), with H2O-drinking animals exhibiting longer latencies when choosing the incorrect stimulus compared to EtOH-drinking animals. There was also a significant group*sex interaction (GLM: β_group*sex_ = −0.31, p=0.04) on incorrect choice latency, with H2O- drinking females displaying longer latencies than both H2O-drinking males (p=0.05) and their EtOH-drinking counterparts (p=0.02). In summary, the data for early reversal learning suggests there was no longer any effect of prior EtOH experience on initiating trials, as had been previously observed during early discrimination learning, but it is important to note that EtOH-drinking animals were less tentative than the H2O-drinking animals given their faster incorrect choice latencies.

#### Last 500 trials

An analysis of initiation latencies and omissions, choice (correct and incorrect) latencies and omissions, and reward latencies was also conducted for the last 500 trials of reversal learning, to capture late phase learning. Animals averaged 155.11±4.06 committed trials per day. Our analyses on initiation omissions indicated females exhibited more initiation omissions (GLM: β_sex_ = −67.96, p=0.02; **Fig 5E**), but there was no effect of group (GLM: β_group_ = −8.46, p=0.85; **Fig 5E**), or group*sex interaction (GLM: β_group*sex_ = −8.79, p=0.85; **Fig 5E**). There was a marginally-significant effect of sex (GLM: β_sex_ = −2.08, p=0.05; **Figs 5F and S5G**), with females taking longer to initiate trials, but no significant group differences (GLM: β_group_ = −0.78, p=0.47; **Figs 5F and S5C**), or group*sex interaction (GLM: β_group*sex_ =0.01, p=0.99; **Fig 5F**). Females also exhibited more choice omissions than males (GLM: β_sex_ = −0.33, p=0.02), but no group differences or group*sex interaction were found. Upon removal of a single outlier, the effect of sex remained (GLM: β_sex_ = −0.33, p=0.01). Males displayed longer correct choice latencies than females (GLM: β_sex_ = 25.46, p=0.003), whereas H2O-drinking animals exhibited longer reward latencies than EtOH-drinking animals (GLM: β_group_ = 0.27, p=0.01; Fig **S5D**). We did not find any other significant effect of group, sex, or group*sex interaction.

Thus, the pattern of prior EtOH experience rendering animals more likely to fail to initiate trials and taking longer to do so in early discrimination learning, was not preserved through reversal learning. In summary, the late reversal phase was characterized by predominantly female-specific attenuations in initiation of trials (both omissions and latencies), as well as correct choice latencies.

### Win-Stay/Lose-Shift (WSLS) Strategies

Potential differences in win-stay/lose-shift (WSLS) strategies on stimulus responses employed by each group (EtOH, H2O) and by sex (male, female) were tested for the first 500 committed trials and last 500 committed trials of the discrimination phase and reversal phase by calculating the frequency of each strategy individually. We compared the frequency of using advantageous strategies (i.e., win-stay, lose-shift) vs. less advantageous strategies (i.e., win-shift, lose-stay) by generating an adaptive score: (win-stay + lose-shift) - (win-shift + lose-stay).

There were no significant effects of group, sex, or a group*sex interaction in stimulusbased strategies individually, or as an ‘adaptive score’ comparison during early or late discrimination learning (**Figs 6A and 6B**). However, during early reversal learning, we found a greater use of the lose-shift strategy in the EtOH-drinking rats than the H20-drinking rats (GLM: β_group_ = −0.10, p=0.01; **Fig S2B**), but a greater lose-stay strategy (GLM: β_group_ =0.10, p=0.01; **Fig S2D**), as well as greater use of the less adaptive strategies (GLM: β_group_ = −0.20, p=0.0002; **Fig 6C**) among the H2O-drinking animals compared to the EtOH-drinking animals. For late reversal learning, we found group differences for all four stimulus-based strategies, with the H2O-drinking animals using the win-stay strategy (GLM: β_group_ = 0.10, p=0.03; **Fig 7A**) and the lose-stay strategy (GLM: β_group_ =0.05, p=0.04; **Fig 7D**) more than the EtOH-drinking animals, suggesting overall more stimulus persistence. Conversely, the EtOH-drinking animals used the win-shift strategy (GLM: β_group_ = −0.10, p=0.03; **Fig 7C**) and lose-shift strategies (GLM: β_group_ = −0.05, p=0.04; **Fig 7B**) more than the H2O-drinking animals, indicating more of an exploration-based strategy. No significant effects were uncovered for adaptive score in the late reversal phase.

**Fig 6.**
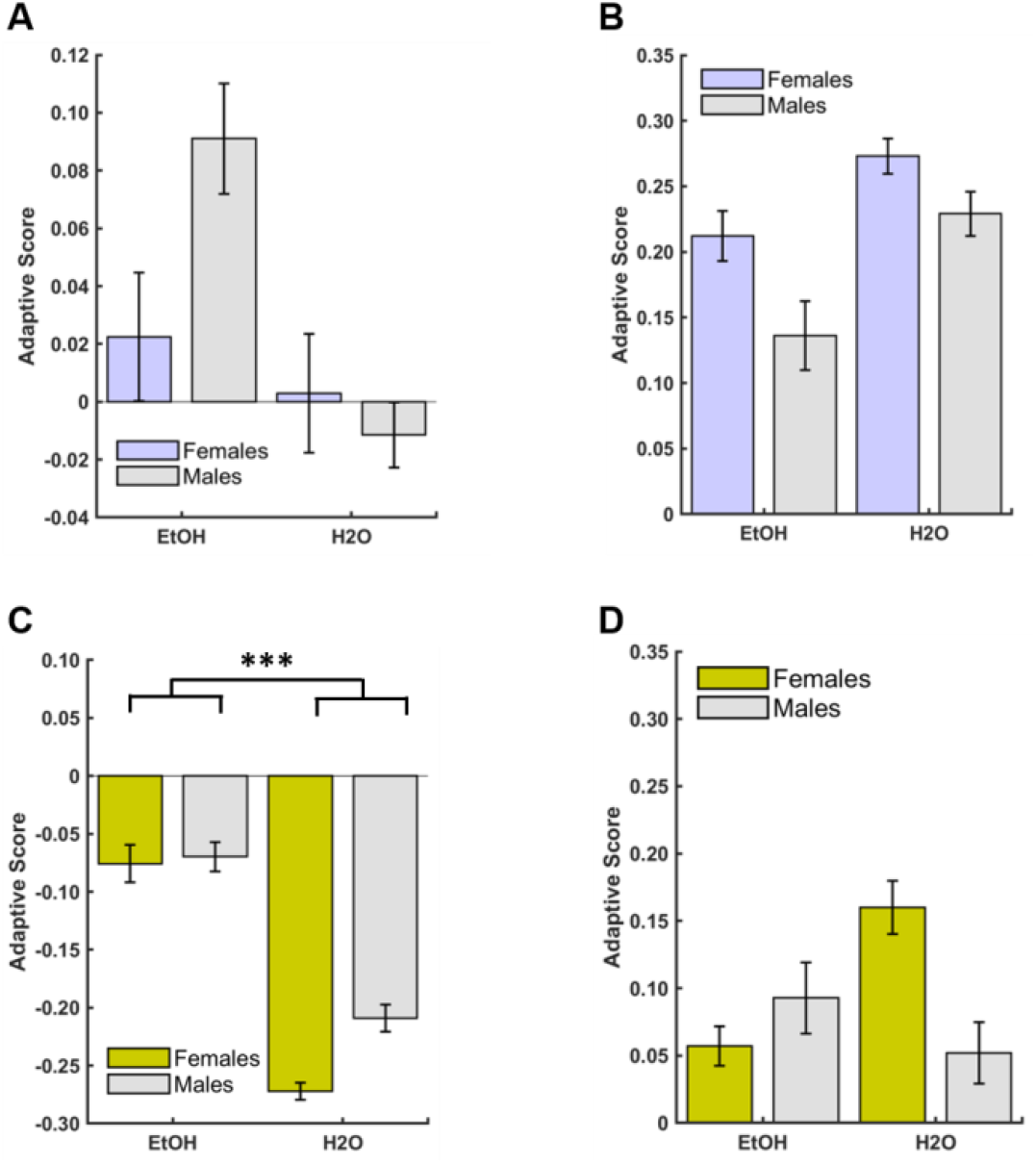
Drinking group differences in early reversal learning strategies. An adaptive score was calculated as the difference between advantageous strategies and less advantageous strategies: (win-stay + lose-shift) - (win-shift + lose-stay). (**A**) There were no group or sex differences in adaptive scores during early discrimination learning. (**B**) There were no group or sex differences in adaptive scores during late discrimination learning. (**C**) EtOH group exhibited higher adaptive scores than H2O group during early reversal learning, but there were no sex differences. (**D**) There were no group or sex differences in adaptive scores during late reversal learning. Bars indicate ± S. *E. M.* n=16 males, n=16 females, *p ≤0.05, **p ≤0.01 ***p ≤0.001

**Fig 7.**
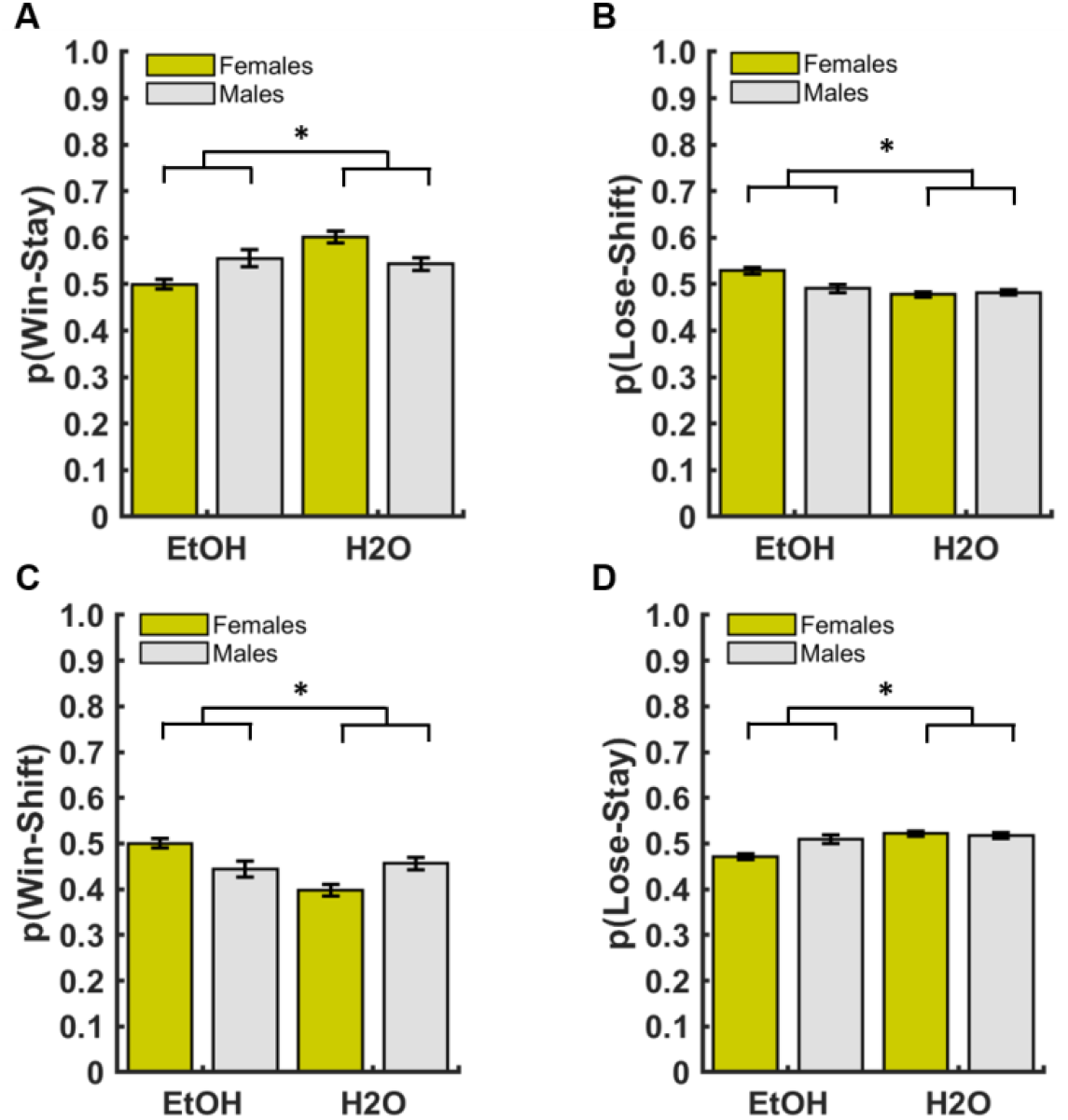
Drinking group differences in the use of Win-Stay/Lose-Shift strategies during late probabilistic reversal learning. (**A**) H2O-drinking animals used the win-stay strategy more than EtOH-drinking animals. (**B**) EtOH-drinking animals used the lose-shift strategy more than H2O-drinking animals. (**C**) EtOH-drinking animals used the win-shift strategy more than H2O-drinking animals. (**D**) H2O-drinking animals used the lose-stay strategy more than EtOH-drinking animals. Bars indicate ± S. *E. M.* *p ≤0.05

## Discussion

The present study used an intermittent access model to study the effect of voluntary EtOH consumption on cognitive flexibility using a probabilistic reversal learning paradigm. We included sex as an a-priori moderator. Although forced-exposure models such as intraperitoneal (i.p.) injections (45,46) and EtOH vapor inhalation (47–49) are well-established methods in rodents, they may not be as representative of human alcohol consumption. Therefore, we used a two-bottle choice procedure that allows for oral consumption of EtOH, resulting in increased ecological validity and variability in consumption patterns, which may be important in generating individual differences in alcohol consumption to study subsequent flexible reward learning. To our knowledge, there had only been one study that previously used this voluntary consumption model to test the effects of EtOH on reversal learning, and found no effect (6). However, it is important to note that although the rats in that EtOH group had access to EtOH for 6 weeks, the rats in that study did not demonstrate escalation normally seen with intermittent voluntary EtOH consumption models, including our study. Though we also corroborate no pronounced effects of EtOH exposure on overall learning, a more fine-grained analysis of trial-by-trial and latency data revealed that EtOH-experienced animals were less likely to initiate trials and were slower to initiate trials throughout pretraining and discrimination learning. Since EtOH drinking rats were unaffected in their reward collection and stimulus response times, collectively the data support the interpretation that the most pronounced attentional decrements appearing closest in time to drinking, despite intact motivation for reward and motor responding throughout learning. We further elaborate on these attentional effects, as well as reversal-specific EtOH effects on WSLS strategies below.

### Consumption patterns of EtOH

We observed an escalation of drinking over the course of the twenty-nine days of alcohol access, irrespective of sex, an expected pattern when using intermittent-access models compared to continuous access models. Several studies administering intermittent exposure have shown that alternating brief periods of alcohol access with brief periods of no access can actually escalate alcohol consumption to excessive levels (25,50–55) compared to continuous daily access (25,50,51) which typically exhibit more moderate, but stable levels of intake. However, despite the recent popularity of the intermittent-access model, the underlying psychological and neurobiological mechanisms that promote the escalation of alcohol consumption remain unclear and should be investigated in future studies.

We found that females reached a higher EtOH consumption level and exhibited greater high-drinking days than males. These findings are consistent with previous studies showing that female rodents drink more EtOH than males (22–25) and exhibit less aversion to EtOH, as demonstrated by conditioned taste aversion using EtOH-saccharin pairings (27–29), with males developing an aversion after only one pairing and females after the third pairing and only at higher doses of EtOH (28). Other groups have reported that the rewarding effects of EtOH are enhanced in females and therefore, may be hormone-dependent (56), which may explain the increased EtOH intake over time that may lead to increases in potential for over-consumption. However, the role of gonadal hormones on ethanol intake and preference remain unclear, as other studies have shown that the removal of testicular hormones in males decreases alcohol intake, and no differential consumption in ovariectomized vs. intact females (24). Although seemingly contradictory, these findings may provide further evidence of the dissociation between chromosomal and gonadal sex, given that studies have found alcohol reinforcement is mediated by chromosomal sex, independent of gonadal phenotype (57). Taken together, the present findings add to a growing body of evidence for sex differences in alcohol consumption patterns.

### Attentional deficits following EtOH across learning stages

We observed the most pronounced impairment following EtOH on sessions to reach criterion during pretraining; a pattern that was not maintained through discrimination or reversal learning. Prior studies testing the relationship between alcohol exposure and performance on reversal learning tasks have largely been mixed, with some studies demonstrating alcohol produced impairments in both discrimination and reversal learning (4,58), other showing no impairments for either (5,6,45,59,60), and some only showing impairments on reversal, with the discrimination learning phase largely intact (5,47,58,61,62). These conflicting findings may be due to variations in alcohol administration procedures, most of which have used forced-exposure models, variations in maximum blood ethanol concentration (BEC) levels, and/or types of reversal learning paradigms employed. Even similar methods of administrations produce variable results with studies showing vapor EtOH exposure impairs reversal learning (47), does not impair (60), or improves reversal learning (59), with the doses of i.p. EtOH administration determining whether an impairment is observed (45). Our study is consistent with the only other study to our knowledge that has used a 2-bottle choice procedure to assess effects on reversal learning, which similarly found no overt learning impairment (6). Although we did not measure BEC levels during the 2- bottle choice procedure, it is likely they never reached BECs known to impair reversal learning based on previous experiments using forced-exposure models (150-550 mg/dl). Task parameters (i.e. stimulus modalities, probability of reward) may also contribute to differential effects, with groups using lever or touchscreen-based responding reporting no pronounced impairments on discrimination or reversal learning (5,6,59), whereas groups employing Morris Water and Barnes Maze tasks reporting impairments in reversal learning. Discrimination learning seems to remain mostly intact across diverse paradigms (58,61–63). We maintain that many past studies of EtOH effects on reversal learning typically report omnibus measures of learning and do not probe more micro (trial-by-trial) analyses that may be more sensitive to EtOH effects, as reported here.

We found impairments associated with prior EtOH experience, such that animals previously exposed to EtOH required more sessions to reach criterion and exhibited longer initiation and choice latencies during pretraining. Similar deficits were also found during discrimination learning, with a greater number of initiation omissions and longer initiation latencies in the EtOH-experienced group. These effects were most pronounced in female animals (discussed below). There have been studies of EtOH exposure on attention using the 5-choice serial reaction time task (64–72), considered to be the gold standard for measuring attention in rodents, revealing that EtOH-exposed animals exhibit attention deficits. An evaluation of attentional capacity using a 5-choice continuous performance task following EtOH exposure in rats found this group exhibited more omissions and longer choice latencies relative to control rats, while motivation remained intact. Indeed, there were no differences in accuracy or reward latencies (66), similar to our present findings. However, it is important to note these differences were observed only during acute, not prolonged, abstinence from EtOH exposure- the latter, as we report here. Other groups have previously reported an EtOH dose-dependent decrease in the ability to direct and sustain attention to brief stimuli, but not a complete disruption in overall performance (i.e. percentage correct), also suggestive of an impairment in attentional processing (70). Similarly, we found differences in measures of attention processing (i.e. initiation omissions and latencies), but observed no overall performance deficit in the probability of choosing the better option for both the discrimination and reversal phases of learning. Importantly, attentional deficits following EtOH experience have also been found in human binge-drinkers (i.e. more omitted trials, lower accuracy), particularly under task variants meant to increase attentional load in a human version of the 5-CSRTT (64). Indeed, the pattern we observed here- that EtOH-drinking animals exhibited more initiation omissions and longer initiation latencies (particularly in early phases of pretraining and discrimination)- stand in contrast to their quick reward collection times and intact accuracy measures, relative to H2O-drinking animals in these same phases.

### Sex differences in pretraining and reversal learning

Some interesting sex differences on reversal learning emerged in our experiment. The most pronounced impairments were observed during late reversal, with females exhibiting greater omissions (initiation and choice) and longer initiation latencies than males, irrespective of prior EtOH exposure. This is in agreement with the human literature, which has shown that males outperform females on reversal learning (73,74), and with observations in marmosets where females require more trials to learn reversals than males. Interestingly, though we find sex differences in reversal learning, there were no differences in the number of omitted trials or reaction times (i.e. latencies) (75). Relevant to this, Grissom et al. (2019) conducted an extensive review of sex differences in several aspects of executive function, including attention, and did not find evidence to support robust sex differences in this domain. Prior studies have reported that male rodents show higher levels of novelty-seeking (76), with higher novelty-seeking related to higher levels of impulsivity in males relative to females (77).

### Sex-dependent EtOH effects in early learning and attentional measures

A sex-dependent drinking group difference was observed, with EtOH-exposed females more affected than males on measures of attention: they exhibited more initiation omissions than their H2O-drinking counterparts during both pretraining and early discrimination learning, which is also reflected in a greater number of sessions required to reach criterion in early learning. Although prior research has not provided sufficient evidence supporting an attentional deficit specific to females (78), there is now substantial evidence to support a potential EtOH-specific effect on attentional processing (64,66,70). Therefore, it is plausible that sex effects we observe here are moderated by EtOH-experience, resulting in more pronounced deficits in attentional processing in EtOH-drinking females. It is worth noting that EtOH-exposed males also exhibited some impairments (i.e. more sessions to reach criterion and initiation omissions), but this effect was only observed early in pretraining and did not extend to discrimination or reversal learning.

### Sex-independent EtOH effects on WSLS in reversal learning

We observed sex-independent EtOH effects on WSLS strategies during reversal learning. Rats with prior EtOH experience were more likely to use a “shift” strategy whereas H2O-drinking animals were more likely to “stay” with the previous stimulus choice in the reversal phase. Similarly, animals with prior EtOH experience were generally more flexible in early reversal learning (i.e. they exhibited a greater ‘adaptive’ score) than H2O-drinking animals. This suggests that EtOH-experienced rats had a more tenuous representation of stimulus-outcome contingencies upon criterion-level performance than the H2O-drinking control rats, and could consequently be more flexible. However, all rats generally increased their choice of the better option and rewards collected over time, while decreasing the number of initiation omissions for both the discrimination and reversal phase. The lack of pronounced EtOH impairments on overall discrimination and reversal phases of learning- as measured by global measures such as the probability choosing the better option over time- may be attributed to plasticity following protracted abstinence in rodents (79–81), and humans (82,83). Similarly, the probability of using WSLS strategies across time was ~0.5, suggesting these strategies were not used effectively for learning. Indeed, dissociations in learning and WSLS have been reported before (84). It will be important to investigate the extent to which the later pro-exploratory phenotype relies on an early attentional decrement, or if these are orthogonal effects of chronic EtOH experience.

### Conclusions

In summary, we observed pronounced trial initiation omissions following EtOH experience in females during pretraining and discrimination learning. These phases are closest in time to the last EtOH experience and constitute the early abstinence period. Additionally, this attentional decrement, which was most pronounced in female animals, was partnered by an enhanced exploration strategy in all EtOH drinking animals, both males and females, later in reversal learning.

Alterations related to attention and processing speed in early EtOH abstinence (during pretraining) may have a domino effect on later learning, leading to the sex by drinking group interaction we observe in discrimination learning, and perhaps contribute to the enhanced exploration phenotype in reversal learning. A true test of this would require animals to undergo pretraining, discrimination learning, and drinking prior to any reversal learning. Ultimately, all rats exhibited intact motivation and motor timing, and were able to increase their probability of choosing the better option and number of rewards, while decreasing their initiation omissions. Although voluntary alcohol consumption models, such as the one employed here, do not model severe alcohol dependence like forced-exposure models, they do however reflect escalating, chronic intermittent drinking that corresponds to the early stages of problematic drinking, before individuals transition to alcohol dependence. Attenuated attentional mechanisms in early abstinence may not contribute to decrements in flexible learning per se, but may instead detract from executive functions important in limiting (over)consumption. Future studies should investigate the brain mechanisms and the role of gonadal hormones on alcohol consumption and attention, and systematically compare these measures as predictors of consumption (i.e. relapse) during acute vs. prolonged abstinence.

## Funding

This work was supported by UCLA’s Division of Life Sciences Recruitment and Retention fund (Izquierdo), R01 DA047870 (Izquierdo), and R01 AA024527 (Spigelman). We acknowledge UCLA’s Graduate Division Graduate Summer Research Mentorship and Graduate Research Mentorship programs (Aguirre) and the NSF Graduate Research Fellowship (Aguirre). We also acknowledge Nancy M. Biram Research Fund in Life Sciences Award (Stolyarova), UCLA’s Dissertation Year Fellowship (Stolyarova) and Charles E. and Sue K. Young Graduate Student Fellowship (Stolyarova).

## Acknowledgements

We thank members of the Izquierdo lab for feedback on earlier versions of this work.

## Author contributions

**Conceptualization:** AI, CA, LR, **Data curation:** AS, CA, VM, **Formal analysis:** CA, AS, AI **Funding acquisition:** AI, IS, **Investigation:** CA, KD, SK, AI, **Methodology:** AI, CA, **Project administration:** CA, AI, **Resources:** AI, IS, **Software:** AS, CA, AI, **Supervision:** CA, AI **Validation:** CA, AI, **Visualization:** CA, AI, **Writing – original draft**: CA, AI, **Writing – review and editing:** CA, AI, VM, IS, LR

## Supplemental Figures

**Fig S1.**
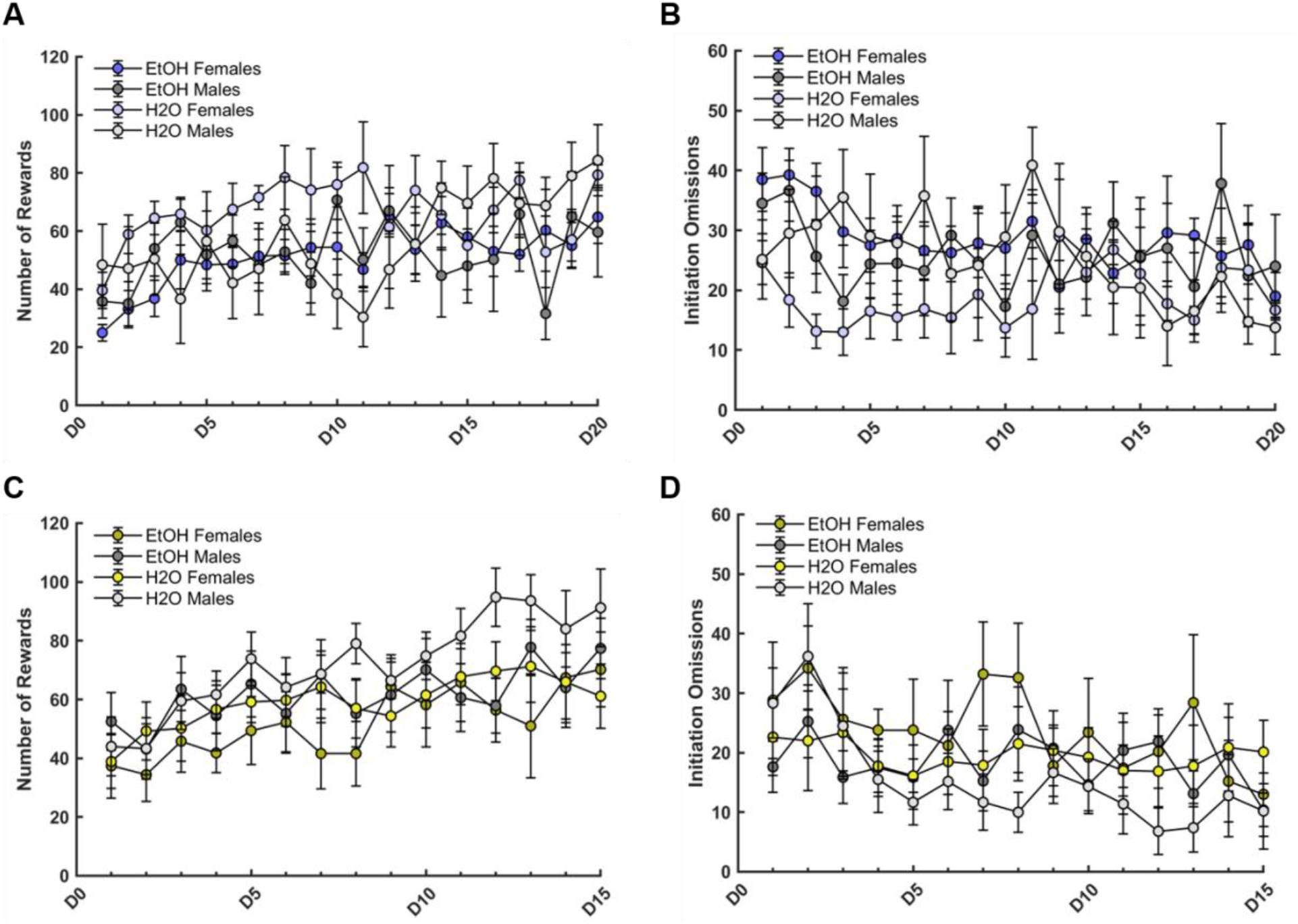
Drinking group and sex differences on number of rewards and initiation omissions during probabilistic discrimination and reversal learning. (A) All animals regardless of drinking group or sex increased their number of rewards collected over the twenty testing days of discrimination learning. H2O-drinking animals and males collected a greater number of rewards than EtOH-drinking animals and females, respectively. Females displayed a greater increase in rewards collected across days. (B) All animals regardless of drinking group or sex decreased their initiation omissions over the twenty testing days of discrimination learning. EtOH-drinking animals and females had more initiation omissions than H2O-drinking animals and males, respectively. EtOH-drinking females displayed more initiation omissions than H2O-drinking females. (C) All animals regardless of drinking group or sex increased their number of rewards collected over the fifteen testing days of reversal learning. There were no group or sex differences on number of rewards collected. (D) All animals regardless of drinking group or sex decreased the number of initiation omissions over the fifteen testing days of reversal learning. There were no group or sex differences on initiation omissions. Bars indicate ± S. *E. M.*

**Fig S2.**
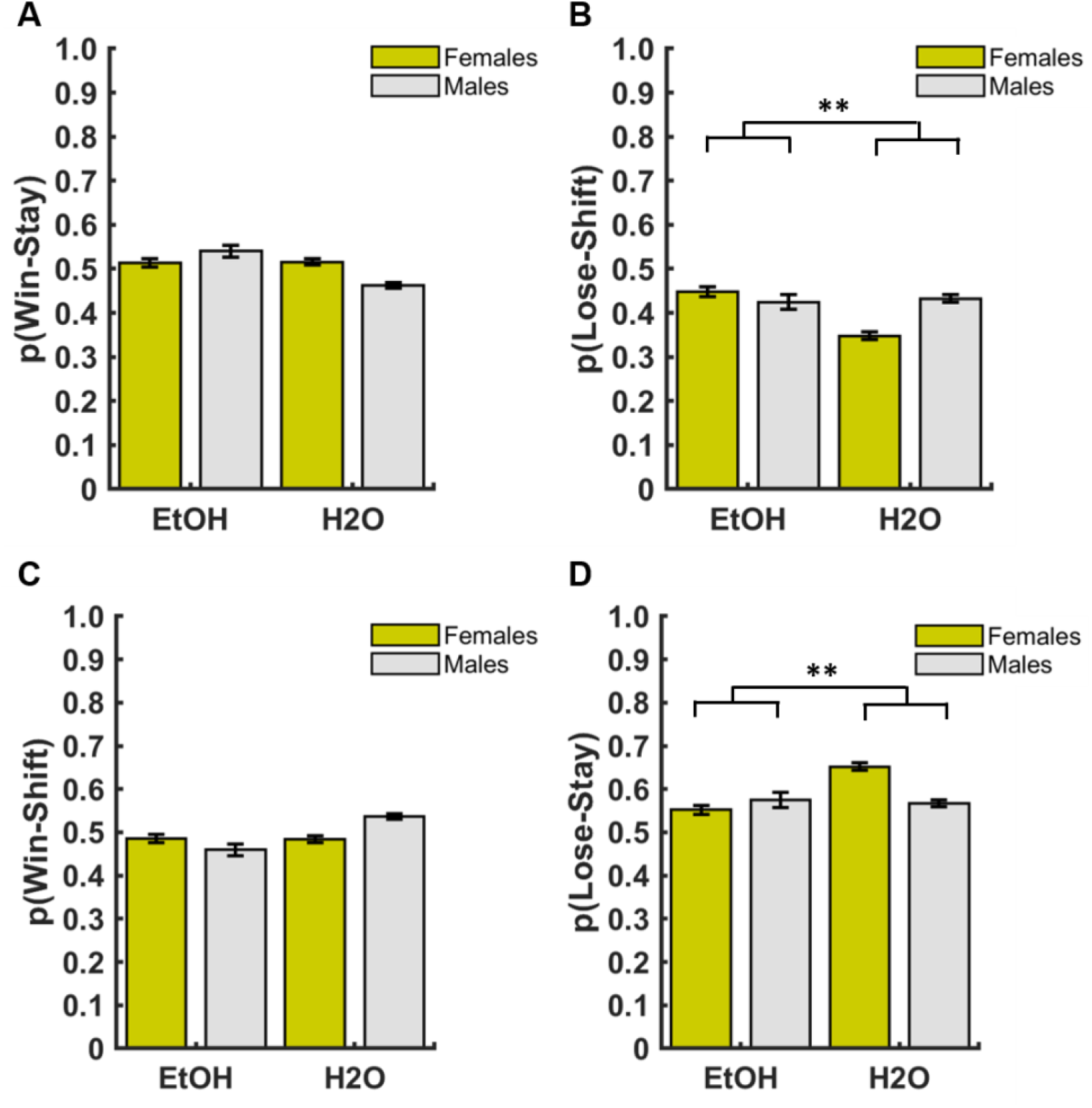
Drinking group differences in use of lose-shift and lose-stay strategies during early probabilistic reversal learning. (A) No group or sex differences in the use of the win-stay strategy. (B) EtOH-drinking animals used the lose-shift strategy more than H2O-drinking animals. (C) No group or sex differences in the use of the win-shift strategy. (D) H2O-drinking animals used the lose-stay strategy more than EtOH-drinking animals. Bars indicate ± *S.E.M.* *p ≤0.05, **p ≤0.01

**Fig S3.**
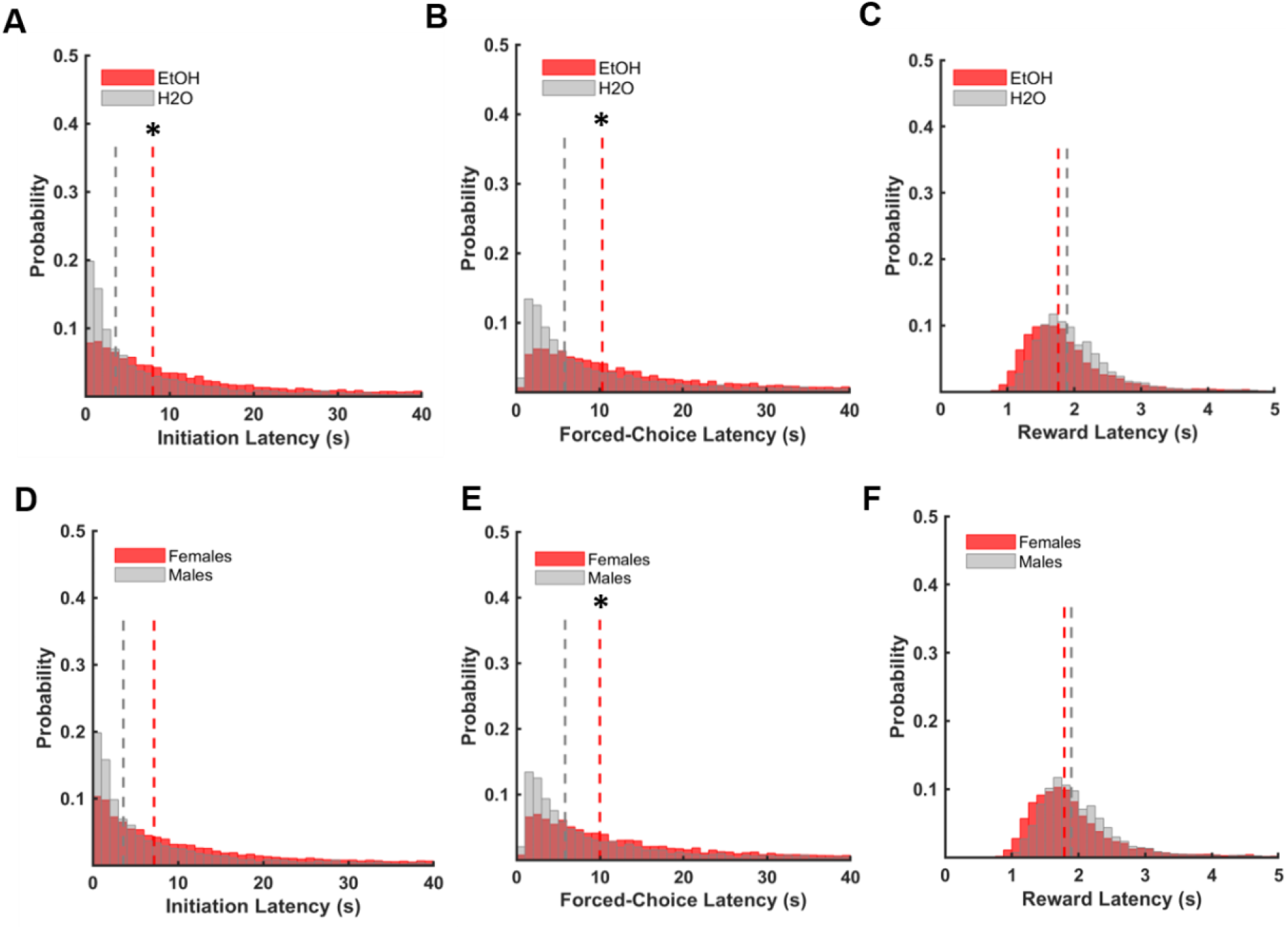
Drinking group and sex differences in latencies during operant pretraining. (A) EtOH group exhibited longer initiation latencies than the H2O group. (B) EtOH group exhibited longer forced-choice latencies than the H2O group. (C) No group differences were found for reward latencies. (D) No sex differences were found for initiation latencies. (E) Females exhibited longer forced-choice latencies than males. (F) No sex differences were found for reward latencies. Dashed lines in latency histograms represent group medians. Bars indicate ± *S.E.M.* n=16 males, n=16 females, *p<0.05

**Fig S4.**
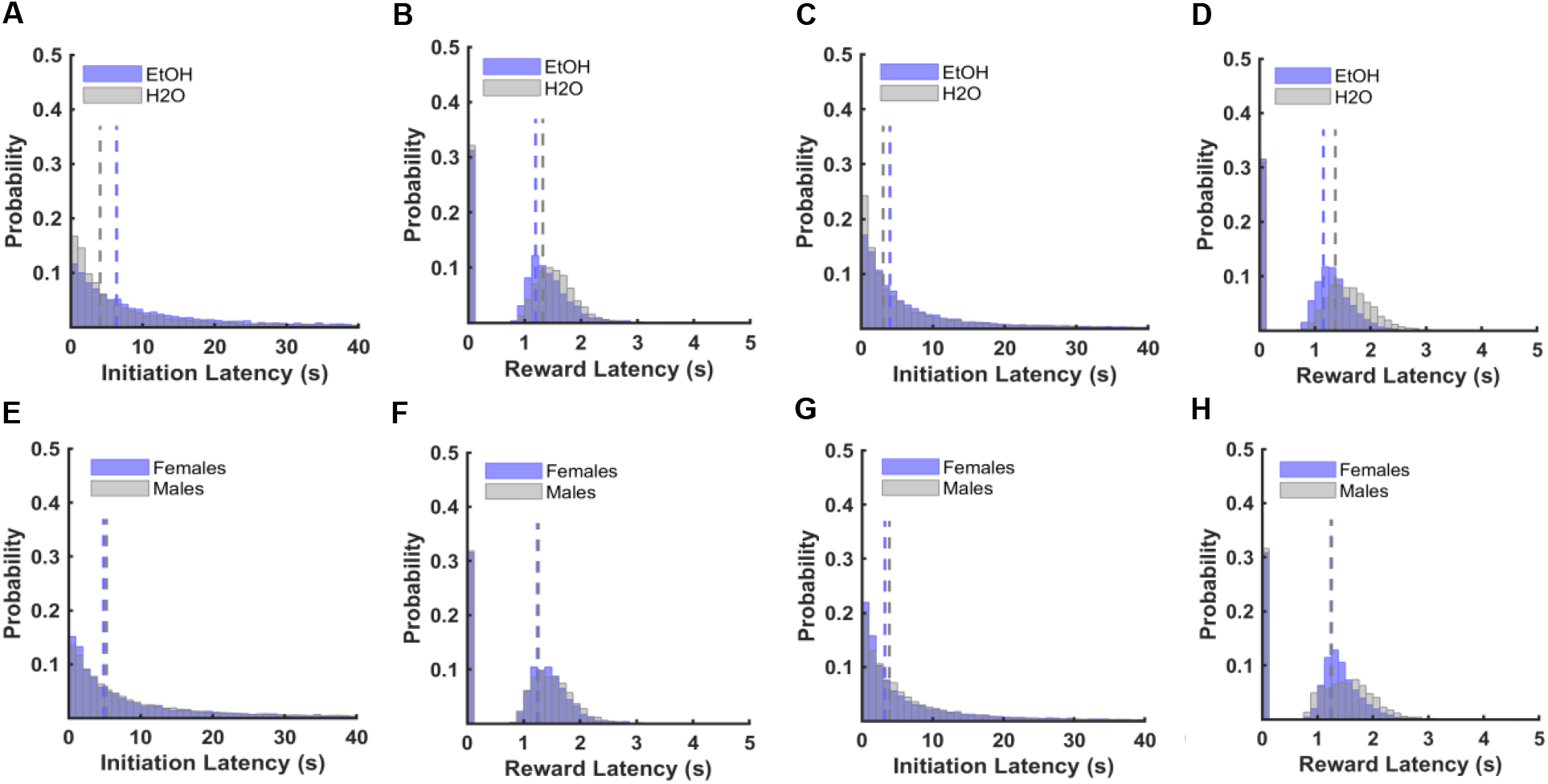
No drinking group or sex differences in latencies during early or late probabilistic discrimination learning. (A) No group differences were found for initiation latencies for the first 500 trials. (B) No group differences were found for reward latencies for the first 500 trials. (C) No group differences were found for initiation latencies for the last 500 trials. (D) No group differences were found for reward latencies for the last 500 trials. (E) No sex differences were found for initiation latencies for the first 500 trials. (F) No sex differences were found for reward latencies for the first 500 trials. (G) No sex differences were found for initiation latencies for the last 500 trials. (H) No sex differences were found for reward latencies for the last 500 trials. Dashed lines in histograms of latencies represent group medians. Bars indicate ± *S.E.M.* n=16 males, n=16 females.

**Fig S5.**
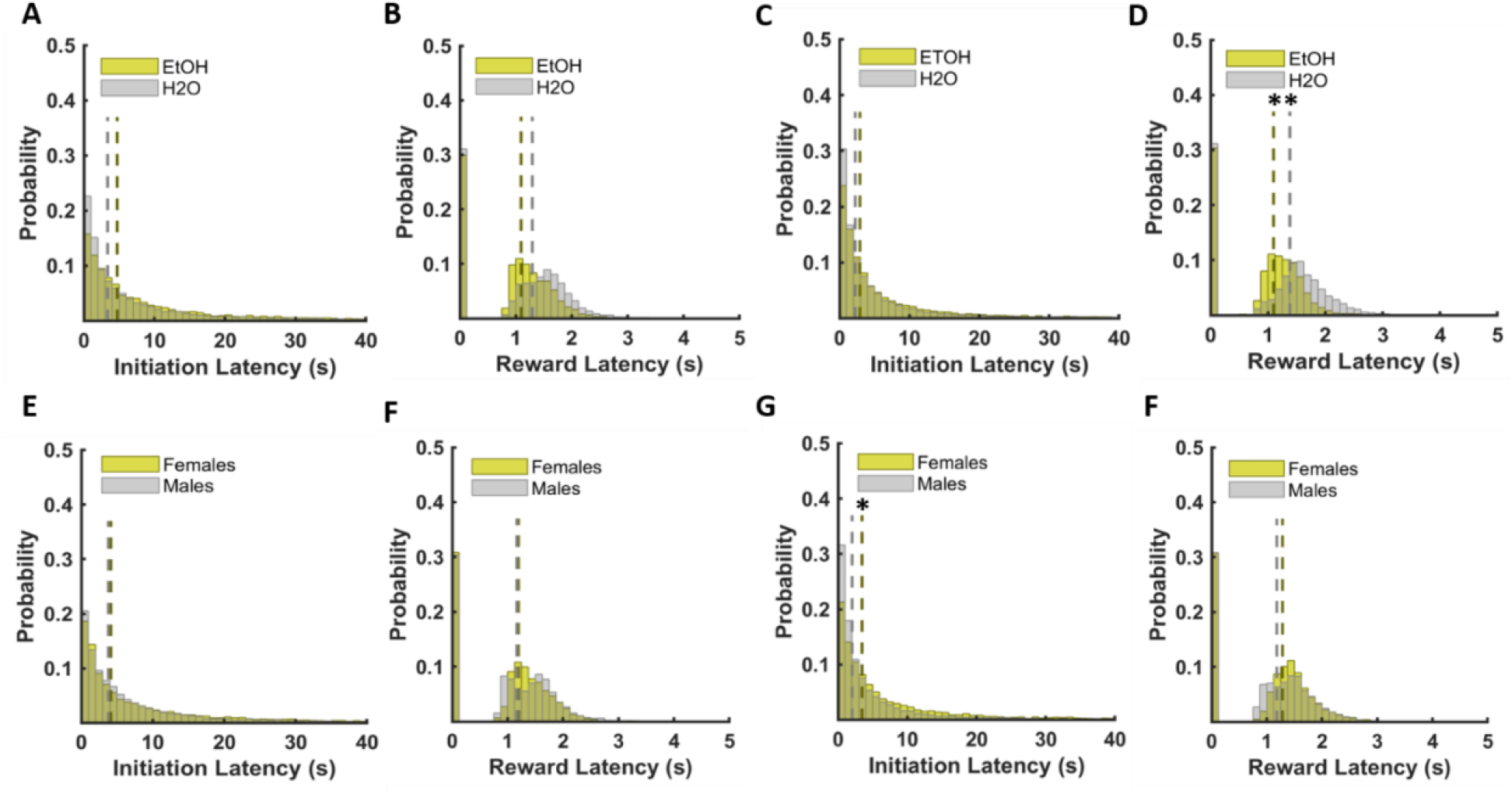
Drinking group and sex differences in latencies during early and late probabilistic reversal learning. (A) No group differences were found for initiation latencies for the first 500 trials. (B) No group differences were found for reward latencies for the first 500 trials. (C) No group differences were found for initiation latencies for the last 500 trials. (D) H2O group exhibited longer reward latencies compared to the EtOH group for the last 500 trials. (E) No sex differences were found for initiation latencies for the first 500 trials. (F) No sex differences were found for reward latencies for the first 500 trials. (G) Females exhibited longer initiation latencies compared to males for the last 500 trials. (H) No sex differences were found for reward latencies for the last 500 trials. Dashed lines in histograms of latencies represent group medians. Bars indicate ± *S.E.M.* n=16 males, n=14 females, *p ≤0.05, **p ≤0.01

## Notes

### Competing Interest Statement

The authors have declared no competing interest.

## References

1. Dajani DR, Uddin LQ. Demystifying cognitive flexibility: Implications for clinical and developmental neuroscience. Trends in neurosciences. 2015 Sep;38(9):571–8.

2. Houston RJ, Derrick J, Leonard K, Testa M, Quigley B, Kubiak A. Effects of Heavy Drinking on Executive Cognitive Functioning in a Community Sample. Addictive behaviors. 2014 Jan;39(1):345–349.

3. Izquierdo A, Brigman JL, Radke AK, Rudebeck PH, Holmes A. The neural basis of reversal learning: An updated perspective. Neuroscience. 2017 Mar 14;345:12–26.

4. Fernandez GM, Stewart WN, Savage LM. Chronic Drinking During Adolescence Predisposes the Adult Rat for Continued Heavy Drinking: Neurotrophin and Behavioral Adaptation after Long-Term, Continuous Ethanol Exposure. Homberg J, editor. PLoS ONE. 2016;11(3):e0149987.

5. Fernandez GM, Lew BJ, Vedder LC, Savage LM. Chronic intermittent ethanol exposure leads to alterations in brain-derived neurotrophic factor within the frontal cortex and impaired behavioral flexibility in both adolescent and adult rats. Neuroscience. 2017 Apr 21;348:324–34.

6. Fisher H, Bright N, Gallo M, Pajser A, Pickens CL. Relationship of low doses of alcohol voluntarily consumed during adolescence and early adulthood with subsequent behavioral flexibility. Behavioural Pharmacology [Internet]. 2017;28(7). Available from: https://journals.lww.com/behaviouralpharm/Fulltext/2017/10000/Relationship_of_low_doses_of_alcohol_voluntarily.4.aspx

7. Amodeo LR, McMurray MS, Roitman JD. Orbitofrontal cortex reflects changes in response-outcome contingencies during probabilistic reversal learning. Neuroscience. 2017 Mar 14;345:27–37.

8. Dalton GL, Wang NY, Phillips AG, Floresco SB. Multifaceted Contributions by Different Regions of the Orbitofrontal and Medial Prefrontal Cortex to Probabilistic Reversal Learning. J Neurosci. 2016 Feb 10;36(6):1996.

9. Ray MH, Hite T, Gallo M, Pickens CL. Operant over-responding is more sensitive than reversal learning for revealing behavioral changes after withdrawal from alcohol consumption. Physiology & Behavior. 2018 Nov 1;196:176–84.

10. McMurray MS, Amodeo LR, Roitman JD. Effects of voluntary alcohol intake on risk preference and behavioral flexibility during rat adolescence. PloS one. 2014 Jul 9;9(7):e100697–e100697.

11. Amitai N, Markou A. Disruption of Performance in the Five-Choice Serial Reaction Time Task Induced By Administration of N-Methyl-D-Aspartate Receptor Antagonists: Relevance to Cognitive Dysfunction in Schizophrenia. Biological Psychiatry. 2010 Jul 1;68(1):5–16.

12. Asinof SK, Paine TA. The 5-choice serial reaction time task: a task of attention and impulse control for rodents. J Vis Exp. 2014 Aug 10;(90):e51574–e51574.

13. Bushnell PJ, Strupp BJ. Chapter 7: Asessing Attention in Rodents. In: Buccafusco JJ, editor. Methods of Behavior Analysis in Neuroscience. 2nd ed. Boca Raton, FL: CRC Press/Taylor & Francis; 2009. p. 119–44.

14. Remmelink E, Chau U, Smit AB, Verhage M, Loos M. A one-week 5-choice serial reaction time task to measure impulsivity and attention in adult and adolescent mice. Sci Rep. 2017 Feb 15;7:42519–42519.

15. Guidi M, Kumar A, Foster TC. Impaired attention and synaptic senescence of the prefrontal cortex involves redox regulation of NMDA receptors. J Neurosci. 2015 Mar 4;35(9):3966–77.

16. Marbach F, Zador AM. A self-initiated two-alternative forced choice paradigm for head-fixed mice. bioRxiv. 2017 Jan 1;073783.

17. Bruinsma B, Terra H, de Kloet SF, Luchicchi A, Timmerman AJ, Remmelink E, et al. An automated home-cage-based 5-choice serial reaction time task for rapid assessment of attention and impulsivity in rats. Psychopharmacology (Berl). 2019 Jul;236(7):2015–26.

18. Martin TJ, Grigg A, Kim SA, Ririe DG, Eisenach JC. Assessment of attention threshold in rats by titration of visual cue duration during the five choice serial reaction time task. Journal of Neuroscience Methods. 2015 Feb 15;241:37–43.

19. Bayless DW, Darling JS, Stout WJ, Daniel JM. Sex differences in attentional processes in adult rats as measured by performance on the 5-choice serial reaction time task. Behavioural Brain Research. 2012 Nov 1;235(1):48–54.

20. Turner KM, Peak J, Burne THJ. Measuring Attention in Rodents: Comparison of a Modified Signal Detection Task and the 5-Choice Serial Reaction Time Task. Front Behav Neurosci. 2016 Jan 14;9:370–370.

21. Worthy DA, Maddox WT. A Comparison Model of Reinforcement-Learning and Win-Stay-Lose-Shift Decision-Making Processes: A Tribute to W.K. Estes. Journal of mathematical psychology. 2014 Apr 1;59:41–9.

22. Vetter-O’Hagen C, Varlinskaya E, Spear L. Sex Differences in Ethanol Intake and Sensitivity to Aversive Effects during Adolescence and Adulthood. Alcohol and Alcoholism. 2009 Nov 1;44(6):547–54.

23. Lourdes de la Torre M, Dolores Escarabajal M, Agüero Á. Sex differences in adult Wistar rats in the voluntary consumption of ethanol after pre-exposure to ethanol-induced flavor avoidance learning. Pharmacology Biochemistry and Behavior. 2015 Oct 1; 137:7–15.

24. Vetter-O’Hagen CS, Spear LP. THE EFFECTS OF GONADECTOMY ON AGE-AND SEX-TYPICAL PATTERNS OF ETHANOL CONSUMPTION IN SPRAGUE-DAWLEY RATS. Alcoholism, clinical and experimental research. 2011 Nov;35(11):2039–49.

25. Hwa LS, Chu A, Levinson SA, Kayyali TM, DeBold JF, Miczek KA. Persistent escalation of alcohol drinking in C57BL/6J mice with intermittent access to 20% ethanol. Alcoholism, clinical and experimental research. 2011 Nov;35(11):1938–47.

26. Wallin-Miller KG, Chesley J, Castrillon J, Wood RI. Sex differences and hormonal modulation of ethanol-enhanced risk taking in rats. Drug and Alcohol Dependence. 2017 May 1;174:137–44.

27. Cailhol S, Mormède P. Conditioned taste aversion and alcohol drinking: strain and gender differences. J Stud Alcohol. 2002 Jan 1;63(1):91–9.

28. Sherrill LK, Berthold C, Koss WA, Juraska JM, Gulley JM. Sex differences in the effects of ethanol pre-exposure during adolescence on ethanol-induced conditioned taste aversion in adult rats. Behavioural Brain Research. 2011 Nov 20;225(1): 104–9.

29. Schramm-Sapyta NL, Francis R, MacDonald A, Keistler C, O’Neill L, Kuhn CM. Effect of sex on ethanol consumption and conditioned taste aversion in adolescent and adult rats. Psychopharmacology. 2014 Apr 1;231(8): 1831–9.

30. Marquardt K, Sigdel R, Brigman JL. Touch-screen visual reversal learning is mediated by value encoding and signal propagation in the orbitofrontal cortex. Neurobiology of learning and memory. 2017 Mar;139:179–88.

31. Stolyarova A, Izquierdo A. Complementary contributions of basolateral amygdala and orbitofrontal cortex to value learning under uncertainty. Schoenbaum G, editor. eLife. 2017;6:e27483.

32. Stolyarova A, Thompson AB, Barrientos RM, Izquierdo A. Reductions in Frontocortical Cytokine Levels are Associated with Long-Lasting Alterations in Reward Valuation after Methamphetamine. Neuropsychopharmacology. 2015 Apr;40(5):1234–42.

33. MATLAB and Statistics Toolbox. Natick, Massachusetts, United States: The Mathworks, Inc.; 2013.

34. IBM SPSS Statistics. Armonk, NY: IBM Corp.; 2017.

35. Morales M, McGinnis MM, McCool BA. Chronic ethanol exposure increases voluntary home cage intake in adult male, but not female, Long–Evans rats. Pharmacology Biochemistry and Behavior. 2015 Dec 1;139:67–76.

36. Priddy BM, Carmack SA, Thomas LC, Vendruscolo JCM, Koob GF, Vendruscolo LF. Sex, strain, and estrous cycle influences on alcohol drinking in rats. Pharmacology Biochemistry and Behavior. 2017 Jan 1; 152:61–7.

37. Piano MR, Carrigan TM, Schwertz DW. Sex differences in ethanol liquid diet consumption in Sprague–Dawley rats. Alcohol. 2005 Feb 1;35(2): 113–8.

38. Leeman RF, Heilig M, Cunningham CL, Stephens DN, Duka T, O’Malley SS. REVIEW: Ethanol consumption: how should we measure it? Achieving consilience between human and animal phenotypes. Addiction Biology. 2010 Feb 9;15(2):109–24.

39. Piantadosi P, Lieberman A, Pickens C, Bergstrom H, Holmes A. A novel multichoice touchscreen paradigm for assessing cognitive flexibility in mice. Learning & Memory. 2019 Jan 1;26:24–30.

40. Bryce CA, Howland JG. Stress facilitates late reversal learning using a touchscreen-based visual discrimination procedure in male Long Evans rats. Behavioural Brain Research. 2015 Feb 1;278:21–8.

41. Stolyarova A, Rakhshan M, Hart EE, O’Dell TJ, Peters MAK, Lau H, et al. Contributions of anterior cingulate cortex and basolateral amygdala to decision confidence and learning under uncertainty. Nature Communications. 2019 Oct 17;10(1):4704.

42. Izquierdo A, Belcher AM, Scott L, Cazares VA, Chen J, O’Dell SJ, et al. Reversal-specific learning impairments after a binge regimen of methamphetamine in rats: possible involvement of striatal dopamine. Neuropsychopharmacology: official publication of the American College of Neuropsychopharmacology. 2010 Jan;35(2):505–14.

43. Izquierdo A. Functional Heterogeneity within Rat Orbitofrontal Cortex in Reward Learning and Decision Making. J Neurosci. 2017 Nov 1;37(44):10529.

44. Robbins TW, Cardinal RN. Computational psychopharmacology: a translational and pragmatic approach. Psychopharmacology. 2019 Aug 1;236(8):2295–305.

45. Badanich KA, Fakih ME, Gurina TS, Roy EK, Hoffman JL, Uruena-Agnes AR, et al. Reversal learning and experimenter-administered chronic intermittent ethanol exposure in male rats. Psychopharmacology. 2016 Oct 1;233(19):3615–26.

46. Knapp DJ, Breese GR. Models of Chronic Alcohol Exposure and Dependence. In: Kobeissy FH, editor. Psychiatric Disorders: Methods and Protocols [Internet]. Totowa, NJ: Humana Press; 2012. p. 205–30. Available from: https://doi.org/10.1007/978-1-61779-458-2_13

47. Badanich KA, Becker HC, Woodward JJ. Effects of chronic intermittent ethanol exposure on orbitofrontal and medial prefrontal cortex-dependent behaviors in mice. Behavioral neuroscience. 2011 Dec;125(6):879–91.

48. Wilhelm Clare J., Hashimoto Joel G., Roberts Melissa L., Bloom Shelley H., Andrew Melissa R., Wiren Kristine M. Astrocyte Dysfunction Induced by Alcohol in Females but Not Males. Brain Pathology. 2015 Jun 19;26(4):433–51.

49. Planeta CS. Animal models of alcohol and drug dependence. Revista Brasileira de Psiquiatria. 2013;35:S140–6.

50. Rosenwasser AM, Fixaris MC, Crabbe JC, Brooks PC, Ascheid S. Escalation of intake under intermittent ethanol access in diverse mouse genotypes. Addiction Biology. 2012 Aug 2;18(3):496–507.

51. Becker HC, Ron D. Animal models of excessive alcohol consumption: Recent advances and future challenges. Alcohol. 2014 May 1;48(3):205–8.

52. Carnicella S, Ron D, Barak S. Intermittent ethanol access schedule in rats as a preclinical model of alcohol abuse. Alcohol (Fayetteville, NY). 2014 May;48(3):243–52.

53. Carnicella S, Amamoto R, Ron D. Excessive alcohol consumption is blocked by glial cell line-derived neurotrophic factor. Alcohol. 2009 Feb 1;43(1):35–43.

54. Simms JA, Steensland P, Medina B, Abernathy KE, Chandler LJ, Wise R, et al. Intermittent Access to 20% Ethanol Induces High Ethanol Consumption in Long–Evans and Wistar Rats. Alcoholism, clinical and experimental research. 2008 Oct;32(10):1816–23.

55. George O, Sanders C, Freiling J, Grigoryan E, Vu S, Allen CD, et al. Recruitment of medial prefrontal cortex neurons during alcohol withdrawal predicts cognitive impairment and excessive alcohol drinking. Proc Natl Acad Sci USA. 2012 Oct 30;109(44):18156.

56. Torres OV, Walker EM, Beas BS, O’Dell LE. Female rats display enhanced rewarding effects of ethanol that are hormone dependent. Alcoholism, clinical and experimental research. 2014 Jan;38(1):108–15.

57. Barker JM, Torregrossa MM, Arnold AP, Taylor JR. Dissociation of genetic and hormonal influences on sex differences in alcoholism-related behaviors. J Neurosci. 2010 Jul 7;30(27):9140–4.

58. Kuzmin A, Liljequist S, Meis J, Chefer V, Shippenberg T, Bakalkin G. Repeated moderate-dose ethanol bouts impair cognitive function in Wistar rats. Addiction Biology. 2012 Jan 1;17(1): 132–40.

59. DePoy L, Daut R, Brigman JL, MacPherson K, Crowley N, Gunduz-Cinar O, et al. Chronic alcohol produces neuroadaptations to prime dorsal striatal learning. Proc Natl Acad Sci USA. 2013 Sep 3; 110(36): 14783.

60. Kroener S, Mulholland PJ, New NN, Gass JT, Becker HC, Chandler LJ. Chronic alcohol exposure alters behavioral and synaptic plasticity of the rodent prefrontal cortex. PLoS One. 2012;7(5):e37541–e37541.

61. Coleman Jr LG, He J, Lee J, Styner M, Crews FT. Adolescent Binge Drinking Alters Adult Brain Neurotransmitter Gene Expression, Behavior, Brain Regional Volumes, and Neurochemistry in Mice. Alcoholism: Clinical and Experimental Research. 2011 Apr 1;35(4):671–88.

62. Obernier JA, White AM, Swartzwelder HS, Crews FT. Cognitive deficits and CNS damage after a 4-day binge ethanol exposure in rats. Pharmacology Biochemistry and Behavior. 2002 Jun 1;72(3):521–32.

63. Coleman LG Jr, Liu W, Oguz I, Styner M, Crews FT. Adolescent binge ethanol treatment alters adult brain regional volumes, cortical extracellular matrix protein and behavioral flexibility. Pharmacology, biochemistry, and behavior. 2014 Jan;116:142–51.

64. Sanchez-Roige S, Baro V, Trick L, Peña-Oliver Y, Stephens DN, Duka T. Exaggerated Waiting Impulsivity Associated with Human Binge Drinking, and High Alcohol Consumption in Mice. Neuropsychopharmacology. 2014 Dec 1;39(13):2919–27.

65. Sanchez-Roige S, Peña-Oliver Y, Ripley TL, Stephens DN. Repeated Ethanol Exposure During Early and Late Adolescence: Double Dissociation of Effects on Waiting and Choice Impulsivity. Alcoholism: Clinical and Experimental Research. 2014 Oct 1;38(10):2579–89.

66. Irimia C, Tuong RN, Quach T, Parsons LH. Impaired Response Inhibition in the Rat 5 Choice Continuous Performance Task during Protracted Abstinence from Chronic Alcohol Consumption. PLOS ONE. 2014 Oct 15;9(10):e109948.

67. Irimia C, Wiskerke J, Natividad LA, Polis IY, Vries TJ, Pattij T, et al. Increased impulsivity in rats as a result of repeated cycles of alcohol intoxication and abstinence. Addiction Biology. 2015;20(2):263–74.

68. Boutros N, Der-Avakian A, Markou A, Semenova S. Effects of early life stress and adolescent ethanol exposure on adult cognitive performance in the 5-choice serial reaction time task in Wistar male rats. Psychopharmacology. 2017 May 1;234(9): 1549–56.

69. Brys I, Pupe S, Bizarro L. Attention, locomotor activity and developmental milestones in rats prenatally exposed to ethanol. International Journal of Developmental Neuroscience. 2014 Nov 1;38:161–8.

70. Givens B. Effect of ethanol on sustained attention in rats. Psychopharmacology. 1997 Jul 1;129(2):135–40.

71. Louth EL, Bignell W, Taylor CL, Bailey CDC. Developmental Ethanol Exposure Leads to Long-Term Deficits in Attention and Its Underlying Prefrontal Circuitry. eneuro. 2016 Sep 1;3(5):ENEURO.0267-16.2016.

72. Slawecki CJ. Two-choice reaction time performance in Sprague–Dawley rats exposed to alcohol during adolescence or adulthood. Behavioural Pharmacology [Internet]. 2006;17(7). Available from: https://journals.lww.com/behaviouralpharm/Fulltext/2006/11000/Two_choice_reaction_time_performance_in.6.aspx

73. Overman WH. Sex differences in early childhood, adolescence, and adulthood on cognitive tasks that rely on orbital prefrontal cortex. Brain and Cognition. 2004 Jun 1;55(1):134–47.

74. Evans KL, Hampson E. Sex differences on prefrontally-dependent cognitive tasks. Brain and Cognition. 2015 Feb 1;93:42–53.

75. LaClair M, Febo M, Nephew B, Gervais NJ, Poirier G, Workman K, et al. Sex Differences in Cognitive Flexibility and Resting Brain Networks in Middle-Aged Marmosets. eNeuro. 2019 Jul 25;6(4):ENEURO.0154-19.2019.

76. Palanza P, Morley-Fletcher S, Laviola G. Novelty seeking in periadolescent mice: sex differences and influence of intrauterine position. Physiology & Behavior. 2001 Jan 1;72(1):255–62.

77. Lukkes JL, Thompson BS, Freund N, Andersen SL. The developmental inter-relationships between activity, novelty preferences, and delay discounting in male and female rats. Developmental Psychobiology. 2016 Mar 1;58(2):231–42.

78. Grissom NM, Reyes TM. Let’s call the whole thing off: evaluating gender and sex differences in executive function. Neuropsychopharmacology. 2019 Jan 1;44(1):86–96.

79. Crews FT, Nixon K. Mechanisms of Neurodegeneration and Regeneration in Alcoholism. Alcohol and Alcoholism. 2008 Oct 21;44(2):115–27.

80. Nixon K, Kim DH, Potts EN, He J, Crews FT. Distinct cell proliferation events during abstinence after alcohol dependence: Microglia proliferation precedes neurogenesis. Neurobiology of Disease. 2008 Aug 1;31(2):218–29.

81. Nixon K, Crews FT. Temporally Specific Burst in Cell Proliferation Increases Hippocampal Neurogenesis in Protracted Abstinence from Alcohol. J Neurosci. 2004 Oct 27;24(43):9714.

82. O’Neill J, Cardenas VA, Meyerhoff DJ. Effects of Abstinence on the Brain: Quantitative Magnetic Resonance Imaging and Magnetic Resonance Spectroscopic Imaging in Chronic Alcohol Abuse. Alcoholism: Clinical and Experimental Research. 2001 Nov 1;25(11): 1673–82.

83. Rosenbloom MJ, Rohlfing T, O’Reilly AW, Sassoon SA, Pfefferbaum A, Sullivan EV. Improvement in memory and static balance with abstinence in alcoholic men and women: Selective relations with change in brain structure. Psychiatry Research: Neuroimaging. 2007 Jul 15;155(2):91–102.

84. Verharen JPH, den Ouden HEM, Adan RAH, Vanderschuren LJMJ. Modulation of value-based decision making behavior by subregions of rat prefrontal cortex. Psychopharmacology 2020 https://doi.org/10.1007/s00213-020-05454-7.

